# The highly repetitive genome of *Myxobolus* sp., a myxozoan parasite of fathead minnows

**DOI:** 10.1101/2024.02.09.579586

**Authors:** Viraj R. Muthye, Aralia Leon Coria, Hongrui Liu, Constance A. M. Finney, Cameron P. Goater, James D. Wasmuth

## Abstract

**Background:** The Myxozoa is a group of at least 2,400 endoparasites within the phylum Cnidaria. All myxozoans have greatly reduced in size and morphology compared to free-living members of the phylum. They are best known for causing disease in economically important fish across the world; for example, *Myxobolus cerebralis* causes whirling disease, which can kill 90% of infected juvenile salmonid fish. In 2017, a new myxozoan species was identified in Alberta. *Myxobolus* sp. causes distinct lesions in fathead minnows, which are ultimately fatal. Here, we sequenced, assembled and analyzed the genome of *Myxobolus* sp. to understand how the parasite interacts with its fish host and identify potential strategies to counter this emerging threat.

**Results:** At 185 Mb, the *Myxobolus* sp. genome is the largest myxozoan genome sequenced so far. This large genome size is, in part, due to the high repetitive content; 68% of the genome was interspersed repeats, with the MULE-MuDR transposon covering 18% of the *Myxobolus* sp. genome. Similar to myxozoan genomes, the *Myxobolus* sp. genome has lost many genes well conserved in other eukaryotes. However, we also identified multiple expansions in gene families—serine proteases, hexokinases, and FLYWCH-domain containing proteins—which suggests their functional importance in the parasite. The mitochondrial genome of *Myxobolus* sp. encodes only five of the thirteen protein-coding genes typically found in animals. We found that the mitochondrial gene *atp6* was transferred to the nucleus and acquired a mitochondria-targeting signal in *Myxobolus* sp.

**Conclusions:** Our study provides valuable insights into myxozoan biology and identify promising avenues for future research. We also propose that *Myxobolus* sp. is promising myxozoan model to explore host-parasite interactions in these parasites.

## 1. Introduction

Myxozoa (phylum Cnidaria) are an enigmatic group of approximately 2,400 species of intracellular parasites, which have drastically simplified their morphology, compared to their free-living relatives: jellyfish, corals, and sea anemones (Alama-Bermejo & Holzer, 2021) (Figure 1A). Myxozoans are found in both freshwater and marine environments, and cycle between invertebrate (definitive) and vertebrate (intermediary) hosts (reviewed in (Eszterbauer et al., 2015)). While myxozoans have been identified in birds (Bartholomew et al., 2008), amphibians (Eiras, 2005), reptiles (Eiras, 2005), and even mammals (Friedrich et al., 2000), they are infamous for causing disease in fish, and pose significant ecological, conservation, and economic threats globally (Alama-Bermejo & Holzer, 2021; Hutchins et al., 2021; James et al., 2021; Koel et al., 2006; Sarker et al., 2015).

**Figure 1.**
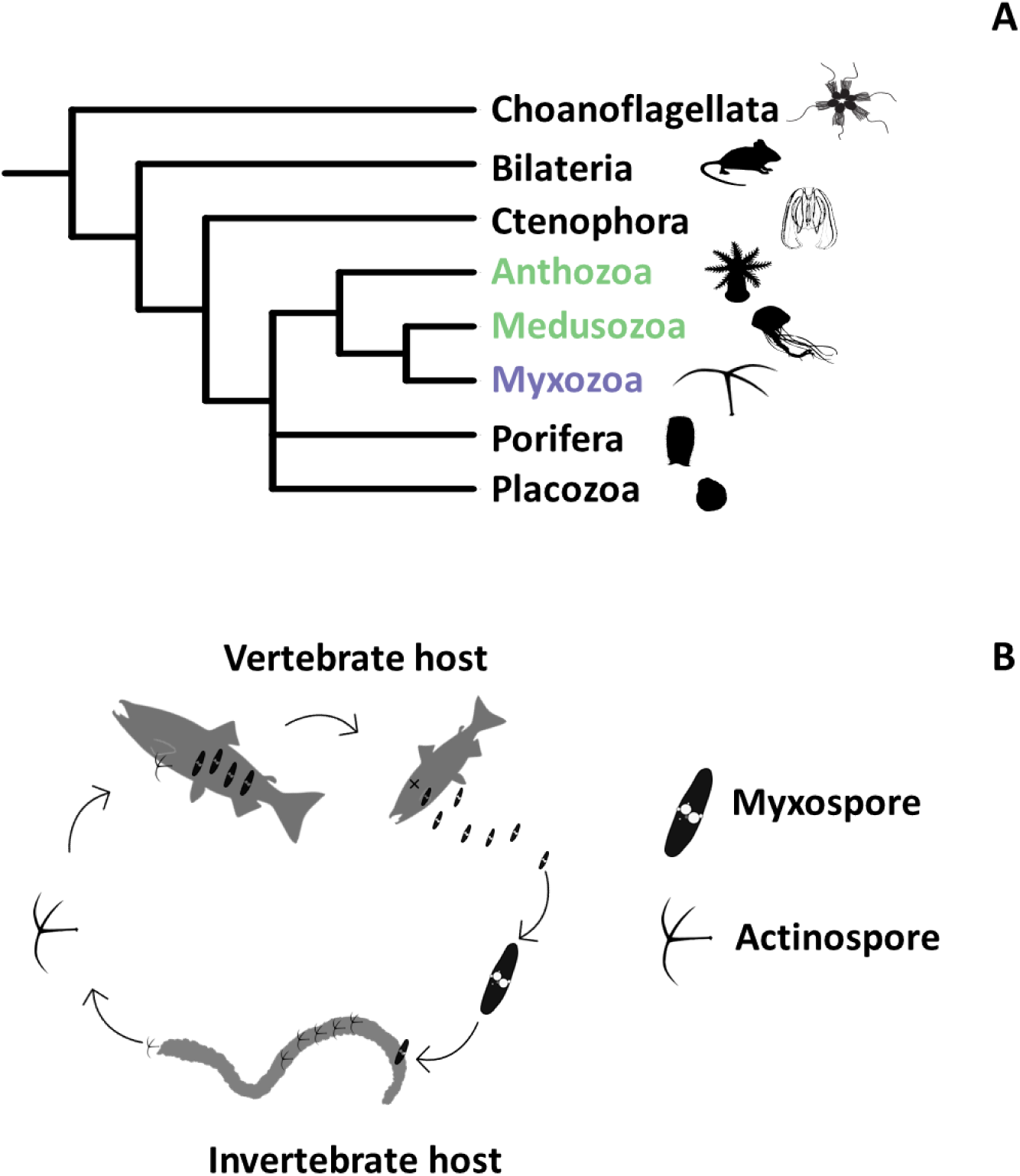
A) Phylogenetic relationships between the five metazoan groups (Porifera, Cnidaria, Ctenophora, Placozoa, and Bilateria) and the outgroup Choanoflagellata, where the free-living classes of phylum Cnidaria are colored in green and Myxozoa is colored in purple. Silhouettes for the animals are taken from PhyloPic (www.phylopic.org). B) A simplified life cycle of *Myxobolus sp.*.

*Myxobolus* (Bütschli 1882) is the most species-rich genus in Myxozoa, with approximately 850 species described so far (Liu et al., 2019). Arguably the most well-known member of the genus is *Myxobolus cerebralis* (Hofer 1903), the causative agent of whirling disease in salmonid fish (Bartholomew & Reno, 2002). *M. cerebralis* attacks the cartilage of juvenile rainbow trout (*Oncorhynchus mykiss*), leading to skeletal deformities. Its definitive host is the oligochaete *Tubifex tubifex* (sludge worm). In 2016, whirling disease was first detected in Canada, in Johnson Lake in Banff National Park, Alberta (Canadian Food Inspection Agency, 2019), and *M. cerebralis* has now been found in four major Albertan water sheds, with approximately 85% of wild rainbow trout fingerlings in the Crowsnest River showing pathological signs of disease (James et al., 2021). This poses severe social, economic, and conservation concerns. The Crowsnest River rainbow trout are sympatric with the endangered westslope cutthroat trout, *Oncorhynchus clarkii lewisi*, which are also susceptible to *M. cerebralis*, raising concerns for their conservation (Koel et al., 2006). Further, rainbow trout is the third most valuable recreational fish species in Alberta, a sport that generates more than $1.1 billion/year for Alberta (Veillard & Clayton, 2020). Multiple other members of the genus cause disease in fish, for example *Myxobolus honghuensis* which infects the pharynx of the economically important gibel carp (*Carassius auratus gibelio*) and *Myxobolus holzerae,* which causes severe gill disease in an Indian major carp (*Labeo rohita*) (Liu et al., 2012; Gupta & Kaur, 2017).

In 2017, a potential new member of the genus, *Myxobolus* sp., was discovered in fathead minnows (*Pimephales promelas*) in southern Alberta (Tilley *et al*. submitted, 2024). As with *M. cerebralis, Myxobolus* sp. uses *T. tubifex* as a definitive host (Figure 1B). Fathead minnows infected with *Myxobolus sp.* have large lesions in the head, ocular, oral, and sinus cavities, leading to reduced olfactory acuity, and increased fatigue, hyperactivity and death compared to uninfected minnows. It is too soon to know the population-level effect on fathead minnows. A concern is that *Myxobolus sp.* could—or already has—spread to other sympatric cyprinid species, including the western silvery minnow (*Hybognathus argyritis*), which is already designated as threatened in Canada (COSEWIC Assessment and Status Report, 2017). Further, fathead minnows have their own important place in the aquatic ecosystem, as a food source for larger fish, and feeding on algae thereby limiting algal overgrowth.

Despite their environmental and economic importance, myxozoans remain under studied. The diagnoses, monitoring, and treatment of parasitic diseases has greatly benefited from knowledge of the genome—and other omics data—for the causative organisms. Myxozoans have been relatively slow to enter the genomic age. To date, genomes have been published for only eight myxozoan species, with transcriptomes available for a further eight species (reviewed in (Alama-Bermejo & Holzer, 2021)). These resources are assisting myxozoan researchers answer questions related to virulence and diversity. Further, of the ∼2400 myxozoan species, only five have a laboratory-confirmed life cycles with just another fifty having partially describe life cycles (Eszterbauer et al., 2015). This knowledge gap has limited efforts to explore myxozoan-host interactions and develop strategies for myxozoan control.

In this study, we set out to sequence, assemble, and annotate the genome of *Myxobolus sp.*. This is only the second time that long-read sequencing technology has been used for the myxozoans, the first being *Myxobolus honghuensis* (Guo et al., 2022). The assembled genome is the largest of any myxozoan sequenced to date. The use of long-read sequencing allowed us to explore the repeat landscape of the parasite genome which showed that *Myxobolus sp.* has one of the most repetitive animal genomes. We compared the genome with other myxozoan and free-living cnidarians and identified multiple gene families that have undergone expansion in *Myxobolus sp.*, like serpins, peroxiredoxins, and hexokinases. In addition, we also present the mitochondrial genome of *Myxobolus sp.* and show that the parasite possesses a highly reduced mitochondrial genome and mitochondrial proteome.

Note: In a previous version of this preprint, we used a species epithet that is under peer-review (February 2024). Our inclusion of it in the preprint was not intended for the purposes of zoological nomenclature. We have removed it in this current version.

## 2. Methods and Materials

### 2.1. Sample Collection, DNA extraction, and Genome Sequencing

*Myxobolus* sp. myxospores were isolated from lesions in fathead minnows following the protocols established by Dr. Cameron Goater and Molly Tilley (Tilley *et al*. 2024 *submitted*). High molecular weight was extracted using the MagAttract® HMW DNA Qiagen kit (Ref.67563). Clean-up used a combination of two kits: the beads from the Select-a-size Magbead kit (Zymo Cat# D4084-3-10) and the buffers from Genomic DNA Clean & Concentrator -25 (Zymo Cat# D4064). A detailed protocol is attached as Supplementary File S1. For sequencing, the SQK-LSK112 Ligation Sequencing Kit (Q20+) from Oxford Nanopore Technologies (ONT) was used to generate the libraries. No changes were made to the steps outlined in the manual for the SQK-LSK112 Ligation Sequencing Kit. The Mk1C MinION sequencer from ONT was used to sequence the genome and Guppy v2.1.7 was used for base-calling with default options.

### 2.2. Genome assembly and contamination removal

To remove the host (fathead minnow) reads, all the base-called reads were mapped to the fathead minnow draft genome (GCF_016745375.1) using minimap2 v2.24 (default options) (Li, 2018; Martinson et al., 2022). Three assemblies for *Myxobolus sp.* from the unmapped reads using different assembly software:

1. FLYE v2.9 (Kolmogorov et al., 2019) (“flye –nano-hq <READSFILE> -o flye -t 48 -g 50m”),
2. wtdbg2 (Ruan & Li, 2020) (“wtdbg2 -x ont -g 50m -t 14 -i <READSFILE> -fo mras; wtpoa-cns -t 14 -i mras.ctg.lay.gz -fo mras.ctg.fa”),
3. Shasta (Shafin et al., 2020) (“shasta-Linux-0.10.0 –input <READSFILE> --threads 14 – config Nanopore-May2022 –Reads.minReadLength 5000”).

BUSCO v5.4.3 (Eukaryota lineage) and N50 was used to select the preferred genome assembly for subsequent annotation and downstream analysis (Simão et al., 2015). The selected assembly was polished with MEDAKA v1.6.0 (https://www.github.com/nanoporetech/medaka). A further round of decontamination was performed using BlobTools v1.1 (Laetsch & Blaxter, 2017), Uniprot Reference Proteomes (downloaded 01 May 2023) (The UniProt Consortium, 2019), and DIAMOND BLASTX (Buchfink et al., 2021). All assembly contigs assigned as chordate, prokaryote, or viral origin were discarded.

### 2.3. RNA extraction and *de novo* transcriptome assembly

Samples of fish muscles containing myxospores were lysed by adding 200 µl of Trizol followed by manual homogenization in dry ice, for a total of three freeze-thaw cycles in a final volume of 600 µl. The resulting mixture was centrifuged, and the supernatant was collected into a new tube where RNA was extracted using the Direct-zol RNA Miniprep kit (Zymo cat. no. R2050) following the manufacturer’s instructions. The extracted RNA was then cleaned using the RNA Clean and Concentrator-5 kit (Zymo cat. no. R1015) following the manufacturer’s instructions. The quantity and quality of the total RNA were assessed on an Agilent 2200 TapeStation RNA ScreenTape. Libraries for sequencing were prepared using the NEBNext Ultra II Directional RNA Library Prep Kit with PolyA capture following the manufacturer’s instructions. Libraries were multiplexed and paired-end sequenced on Illumina NextSeq 500 for 300 cycles (2 × 150 bp) using a mid-output cell, following the manufacturer’s instructions. FastQC v0.11.9 was used to assess the quality of the RNAseq reads before and after filtering reads (Andrews & others, 2010). Adapter sequences and low-quality reads were identified with Trimmomatic v0.39 and removed (Bolger et al., 2014). A *de novo* transcriptome assembly was generated with SPAdes v3.15.4 using default parameters (Bushmanova et al., 2019).

### 2.4. Identification of repeat content and genome annotation

Interspersed repeats and low complexity DNA sequences in the *Myxobolus sp.* genome were masked using RepeatModeller v2.0.1 (Flynn et al., 2020) and RepeatMasker v4.1.4 (www.repeatmasker.org) with default parameters. This soft-masked genome was used as input for BRAKER v2.1.6 to predict protein-coding genes with protein evidence (Brůna et al., 2021). The OrthoDB v10 metazoa protein database was used as the protein evidence for the genome annotation software BRAKER (Kriventseva et al., 2019). BRAKER was also used to annotate the four publicly available myxozoan genome without genome annotations and to re-annotate the four myxozoan genomes which had publicly-available genome annotations. For this, BRAKER was run using protein evidence (OrthoDB v10 metazoa protein database) (Table S2). BUSCO v5.4.3 was used to analyze the completeness of the myxozoan genomes. The lineage used was ‘eukaryota_odb10’ (Creation date: 2020-09-10, number of genomes: 70, number of BUSCO genes: 255). For myxozoan species for whom BRAKER did not improve existing genome annotations based on BUSCO analysis, previously published genome annotations were used for downstream analysis.

### 2.5. Analysis of the mitochondrial genome and proteome

A database of mitochondrial genomes from four myxozoan species, *Polypodium hydriforme,* two cnidarian species, human, *Caenorhabditis elegans*, *Drosophila melanogaster*, and yeast were used to identify whether *Myxobolus sp.* has a mitochondrial genome (Table 1). BLASTN was used to align these genomes to the *Myxobolus sp.* genome (e-value ≤ 1e^-5^) (Camacho et al., 2009). Two methods were used to identify genes in the mitochondrial genome. First, MITOS2 was used to annotate the contig containing the mitochondrial genome with parameters listed in Table 2 (Bernt et al., 2013). MITOS2 was also used to annotate the five publicly available myxozoan mitochondrial genomes. Second, an HMM model approach was used to identify protein-coding genes in the mitochondrial genome. HMM models for the thirteen protein coding genes found in the majority of animal mitochondrial genomes were downloaded from the eggNOG protein database V5.0.0 (Cantalapiedra et al., 2021). HMMer (hmmer.org) was used to search the mitochondrial genome of *Myxobolus sp.* and the publicly-available myxozoan mitochondrial genomes. PROKSEE was used to visualize the mitochondrial genome (Grant et al., 2023).

**Table 1.** Publicly available mitochondrial genomes used in this analysis to identify the mitochondrial genome in *Myxobolus* sp.

| Species | GenBank ID |
| --- | --- |
| *Polypodium hydriforme* | MN794187.1 (Novosolov et al., 2022) |
| *Myxobolus squamalis* | MK087050.1 (Yahalomi et al., 2020) |
| *Kudoa hexapunctata* | LC009437.1 (Takeuchi et al., 2015) |
| *Kudoa septempunctata* | AB731753.1 (Takeuchi et al., 2015) |
| *Kudoa iwatai* | LC009438.1 (Takeuchi et al., 2015) |
| *Nematostella vectensis* | OW052000.1 (Fletcher & Pereira da Conceicao, Lyndell, n.d.) |
| *Dendronephthya gigantea* | NC_013573.1 (Park Eun-Ji et al., 2010) |
| *Homo sapiens* | NC_012920.1 (Anderson et al., 1981) |
| *Caenorhabditis elegans* | NC_001328.1 (Wolstenholme et al., 1994) |
| *Drosophila melanogaster* | NC_024511.2 (Hoskins et al., 2015) |
| *Saccharomyces cerevisiae* | NC_027264.1 (Wolters et al., 2015) |

**Table 2.** The parameters used in MITOS2 to annotate the mitochondria genome of *Myxobolus sp*.

| Property | Value |
| --- | --- |
| Reference | RefSeq 63 Metazoa |
| Genetic Code | 4 |
| Proteins | TRUE |
| tRNAs | TRUE |
| rRNAs | TRUE |
| OH | TRUE |
| OL | TRUE |
| Circular | TRUE |
| Use Al Arab et al. | FALSE |
| E-value Exponent | 2 |
| Final Maximum Overlap | 50nt |
| Fragment Quality Factor | 100 |
| Standard Code | FALSE |
| Cutoff | 50.00% |
| Clipping Factor | 10 |
| Fragment Overlap | 20.00% |
| Local only | TRUE |
| Sensitive only | FALSE |
| ncRNA overlap: | 50 nt |

Most proteins that function in the mitochondria are encoded in the nuclear genome and then imported into the mitochondria (Meisinger et al., 2008). To identify these proteins, human mitochondrial proteins catalogued in MitoCarta v3 (Rath et al., 2021) were downloaded. BLASTP was used to align these human mitochondrial proteins to the proteins from all myxozoan and free-living cnidarian nuclear genomes (e-value ≤ 1e^-5^).

### 2.6. Genome Analysis

#### 2.6.1. Identifying the function of protein-coding genes

If a gene had multiple transcripts, the longest transcript and associated protein per gene was selected. eggNOG-mapper v2.1.9 (Cantalapiedra et al., 2021) to annotate the predicted proteins using default parameters, except two (percentage identity was changed to 30% and all queries were realigned to the PFAM database instead of importing PFAM annotations from orthologs). The output of eggNOG-mapper was parsed to extract 1) PFAM domains, 2) Gene Ontology (GO) terms, and 3) KEGG pathways mapped to each protein.

#### 2.6.2. Gene family analysis

OrthoFinder v2.5.4 was used, with default parameters, to identify groups of orthologous proteins (OGs) between the predicted protein sets of *Myxobolus sp.*, six publicly-available myxozoan species, and seven cnidarian species (listed in Table 3) (Emms & Kelly, 2019). The sets of predicted proteins for the free-living cnidarian species were downloaded from UniProt (one protein per gene). We omitted two myxozoan species (*K. iwatai* and *S. zaharoni*) as they had a high rate of duplication and therefore OrthoFinder identified considerably lower number of single-copy orthologs.

**Table 3.** Details about the seven publicly-available myxozoan genomes used in this study

| Species | Source | Reference |
| --- | --- | --- |
| *Sphaeromyxa zaharoni* | GCA_001455285.1 | (Chang et al., 2015) |
| *Kudoa iwatai* | GCA_001407235.2 | (Chang et al., 2015) |
| *Enteromyxum leei* | GCA_001455295.2 | (Chang et al., 2015; Yahalomi et al., 2017) |
| *Myxobolus squamalis* | GCA_010108815.1 | (Yahalomi et al., 2020) |
| *Henneguya salminicola* | GCA_009887335.1 | (Yahalomi et al., 2020) |
| *Thelohanellus kitauei* | GCA_000827895.1 | (Yang et al., 2014) |
| *Myxobolus honghuensis* | Harvard Dataverse | (Guo, 2021; Guo et al., 2022) |
| *Ceratonova shasta* | GCA_918559645.1 |  |

The single-copy orthologs identified by OrthoFinder were used to analyze phylogenetic relationships between these 14 species. First, all single-copy orthologs were aligned using MAFFT (“--auto” option) (Katoh et al., 2002) and concatenated into a single alignment using the script “catfasta2phyml.pl”. Next, the alignment was trimmed using trimAI (Capella-Gutiérrez et al., 2009). The resulting alignment was used as input for IQ-TREE v2.2.5 (Nguyen et al., 2015). The appropriate substitution model was identified using ModelFinderPlus (“-MFP” option) (Kalyaanamoorthy et al., 2017) based on the Bayesian information criterion (BIC). Branch supports were assessed with 1,000 standard ultrafast bootstrap approximation (UFBoot) replicates (Hoang et al., 2018; Minh et al., 2013). All phylogenetic trees in this study were visualized using iTOL (Letunic & Bork, 2021). CAFE5 (Computational Analysis of gene Family Evolution) was used to identify rapidly expanding and contracting gene families (p-values < 0.05) using the OrthoFinder results and the IQ-TREE-generated phylogenetic tree (Mendes et al., 2021).

Further, 98 proteomes from the class Actinopterygii were downloaded from UniProt. These were used to identify *Myxobolus sp.* proteins that lack sequence similarity to proteins from Actinopterygii. For this, *Myxobolus sp.* proteins were aligned to a database of proteins from these 98 species (e-value ≤ 1e^-3^).

#### 2.6.3. Search for minicollagens and nematogalectins

Protein sequences for minicollagens and nematogalectins from the study done by (Shpirer et al., 2014) were downloaded and BLASTP was used to align them to the myxozoan genomes (e-value ≤ 1e^-5^).

#### 2.6.4. Identification of proteases/inhibitors

The MEROPS Scan Sequences database was downloaded, which is a non-redundant subset of the MEROPS database that is used for the MEROPS batch BLAST searches (Rawlings et al., 2010). BLASTP was used to align the proteins from all species in this study to the MEROPS Scan database (e-value ≤ 1e^-5^).

#### 2.6.5. Searching for homologs of drug target proteins from ChEMBL

All 9,072 single protein drug targets from ChEMBL 33 (www.ebi.ac.uk/chembl) were downloaded. BLASTP was used to align *Myxobolus* sp. proteins to the database of these ChEMBL targets (e-value ≤ 1e^-10^).

## 3. Results

### 3.1. Genome assembly and repeat content analysis

#### 3.1.1. Genome assembly statistics

We extracted DNA from *Myxobolus sp.* myxospores that were isolated from lesions on the heads of fathead minnows. Approximately 3.2 million sequence reads (11.38 Gb) were generated from a single Oxford Nanopore run and passed base-calling filters. Of these, ∼2.1 million reads (65%) could be mapped to the available genome assembly of the fathead minnow, and so were flagged as contamination and removed. The remaining ∼1.1 million reads were used to assemble the *Myxobolus sp.* genome. To achieve the best possible assembly, we generated assemblies from three algorithms: FLYE, wtdbg2, and Shasta. By reviewing assembly metrics—the most complete BUSCO complement and assembly continuity—we selected the FLYE assembly for the subsequent analysis (Table 4). We used BlobTools to remove further contamination by discarding all contigs that were either mapped to bacteria, eukaryote, or viruses (92 contigs). The final *Myxobolus sp.* genome was comprised of 1,503 contigs (185 Mb with N50 of 407.91 Kb). At 185 Mbp, the *Myxobolus sp.* genome assembly is the largest of any myxozoan species sequenced so far (Table 5).

**Table 4.** Comparison of the *Myxobolus sp.* genome assembled using three genome assemblers (FLYE, wtdbg2, and Shasta) based on their BUSCO scores (Eukaryota lineage in the genome mode) and N50 values.

| Assembler | Genome size (nt) | # Contigs | BUSCO (Eukaryota)<br>% where n=225 | N50 |
| --- | --- | --- | --- | --- |
| FLYE | 190,986,082 | 1,595 | C:32.2 [S:31.4, D:0.8],<br>F:19.2, M:48.6 | 383 kb |
| wtdbg2 | 178,055,605 | 1,282 | C:28.2 [S:27.8, D:0.4],<br>F:18.8, M:53.0 | 331 kb |
| Shasta | 165,736,712 | 3,742 | C:27.5 [S:27.1, D:0.4],<br>F:17.3, M:55.2 | 80 kb |

**Table 5.** Genome statistics of all publicly-available myxozoan genomes and *Myxobolus sp*.

| Species | Platform | # Contigs | Size (Mb) | Contig N50 (Kb) | BUSCO (genome mode, eukaryote dataset) % where n=225 |
| --- | --- | --- | --- | --- | --- |
| *Myxobolus sp.* | Oxford Nanopore | 1503 | 184.9 | 407.9 | C:31.4 [S:31.0, D:0.4], F:19.2, M:49.4 |
| *Myxobolus honghuensis* | PacBio + Illumina HiSeq 2500 | 1118 | 161.1 | 1273.5 | C:32.6 [S:31.8, D:0.8], F:20.8, M:46.6 |
| *Myxobolus squamalis* | Illumina HiSeq 3000 | 37921 | 43.7 | 1.3 | C:26.3 [S:26.3, D:0.0], F:17.3, M:56.4 |
| *Sphaeromyxa zaharoni* | Illumina HiSeq 2000 | 70935 | 173.6 | 4.5 | C:30.6 [S:11.0, D:19.6], F:23.1, M:46.3 |
| *Kudoa iwatai* | Illumina HiSeq 2000 / Illumina Genome Analyzer IIx | 1646 | 31.2 | 39.5 | C:31.8 [S:16.1, D:15.7], F:17.3, M:50.9 |
| *Enteromyxum leei* | Illumina HiSeq 2000 | 69178 | 68.2 | 1.0 | C:22.8 [S:20.4, D:2.4], F:21.6, M:55.6 |
| *Henneguya salminicola* | Illumina HiSeq 3000 | 18332 | 61.4 | 1.2 | C:32.9 [S:32.9, D:0.0], F:20.8, M:46.3 |
| *Thelohanellus kitauei* | 454 GS FLX + Illumina Genome Analyzer II | 15632 | 150.3 | 13.0 | C:32.2 [S:31.4, D:0.8], F:18.0, M:49.8 |
| *Ceratonova shasta* | Illumina HiSeq 2000 | 24903 | 69.8 | 9.3 | C:27.1 [S:25.1, D:2.0], F:8.2, M:64.7 |

#### 3.1.2. Repeat content analysis

Approximately 68% of the genome is comprised of interspersed repeats (Table 6). To confirm that the large repeat content in *Myxobolus sp.* was not an artefact of the genome assembler, we also analyzed the repeat content of the genomes assembled using two other tools-wtdbg2 and Shasta. We found that these two genome assemblies were highly repetitive as well (67.2% for the wtdbg2 assembly and 68.2% for the Shasta assembly). In addition, we compared the types and prevalence of different types of repeats with *Myxobolus sp.*. At the time of this analysis, the *M. honghuensis* assembly was the only other myxozoan built from long-reads, giving us confidence in the assembly of repetitive loci. Using the same tools, we found that just over half the *M. honghuensis* genome (51.0%) was comprised of repetitive regions.

**Table 6.** Results of RepeatModeller and RepeatMasker analysis for genomes of *Myxobolus* sp. and *Myxobolus honghuensis*

| Repeat Elements |  | *Myxobolus* sp. |  |  | *Myxobolus honghuensis* |  |  |
| --- | --- | --- | --- | --- | --- | --- | --- |
|  |  | # elements | length occupied | % of sequence | # elements | length occupied | % of sequence |
| Retroelements |  | 14495 | 17182784 | 9.29 | 1121 | 1469706 | 0.91 |
|  | SINEs | 0 | 0 | 0 | 0 | 0 | 0 |
|  | Penelope | 0 | 0 | 0 | 0 | 0 | 0 |
|  | LINEs | 532 | 258231 | 0.14 | 0 | 0 | 0 |
|  | LTR elements | 13963 | 16924553 | 9.15 | 1121 | 1469706 | 0.91 |
| DNA transposons |  | 112916 | 80978766 | 43.79 | 43041 | 23662456 | 14.69 |
|  | hobo-Activator | 6092 | 6575235 | 3.56 | 18240 | 11145247 | 6.92 |
|  | Tc1-IS630-Pogo | 37716 | 22841177 | 12.35 | 12193 | 4972777 | 3.09 |
|  | En-Spm | 0 | 0 | 0 | 0 | 0 | 0.00 |
|  | MULE-MuDR | 39574 | 33412948 | 18.07 | 6488 | 4008242 | 2.49 |
|  | PiggyBac | 41 | 30174 | 0.02 | 733 | 508393 | 0.32 |
|  | Tourist/Harbinger | 0 | 0 | 0 | 0 | 0 | 0.00 |
|  | Other_(Mirage,P-element, Transib) | 0 | 0 | 0 | 0 | 0 | 0.00 |
| Rolling-circles |  | 2705 | 949873 | 0.51 | 0 | 0 | 0 |
| Unclassified |  | 116134 | 27904502 | 15.09 | 166939 | 57023818 | 35.4 |
| Total interspersed repeats |  |  | 126066052 | 68.17 |  | 82155980 | 51.00 |
| Small RNA |  | 0 | 0 | 0 | 0 | 0 | 0 |
| Satellite |  | 0 | 0 | 0 | 0 | 0 | 0 |
| Simple repeats |  | 18636 | 870644 | 0.47 | 36080 | 1727836 | 1.07 |
| Low complexity |  | 8293 | 422235 | 0.23 | 12949 | 668608 | 0.42 |

The *Myxobolus sp.* transposable element (TE) landscape was dominated by DNA transposons, which occupied 43.8 % of the genome and retrotransposons occupied a further 9.3 % of the genome (Table 6). In contrast, the *M. honghuensis* assembly is comprised of 14.7 % DNA transposons and 1% retrotransposons. Of the *Myxobolus sp.* DNA transposons, MULE-MuDR transposons were the most abundant and were identified in 18.1% of the total genomic sequence. Compared to *Myxobolus sp.,* we found that just 2.5% of the genome was occupied by MULE-MuDR elements. It is important to note that more repeats in *M. honghuensis* were unclassified (35.4%) compared to *Myxobolus sp.* (15.1%).

### 3.2. Genome annotation and analysis

We identified 10,106 protein-coding genes in the *Myxobolus sp.* genome, which is around 65% of the number of genes in the *M. honghuensis* genome. The average intron length, exon length and gene lengths in *Myxobolus sp.* and *M. honghuensis* were similar (Table 7). We reannotated all eight publicly-available myxozoan genomes (with protein evidence) (Table 8). Four of these eight genomes lacked publicly-available annotations and a further two species— *Myxobolus squamalis* and *Thelohanellus kitauei*—received improved annotations, based on their BUSCO scores (Table 8).

**Table 7.** Distribution of intron lengths, exon lengths, and gene lengths for *Myxobolus sp.* and *M. honghuensis,* from BRAKER results

|  | Intron lengths |  | Exon length |  | Gene length |  |
| --- | --- | --- | --- | --- | --- | --- |
|  | *Myxobolus sp.* | *M. honghuensis* | *Myxobolus sp.* | *M. honghuensis* | *Myxobolus sp.* | *M. honghuensis* |
| values | 32,234 | 52,165 | 43,272 | 69,808 | 10,106 | 16,867 |
| min | 5 | 4 | 3 | 3 | 196 | 108 |
| max | 59,944 | 24,000 | 12,697 | 12,542 | 89,608 | 113,913 |
| range | 59,939 | 23,996 | 12,694 | 12,539 | 89,412 | 113,805 |
| sum | 24,066,630 | 36,410,620 | 11,571,580 | 17,172,630 | 33,931,130 | 51,515,720 |
| median | 22 | 28 | 155 | 150 | 1,007 | 1,208 |
| mean | 746.62 | 697.99 | 267.41 | 246.00 | 3,357.52 | 3,054.23 |
| std.dev | 3,262.03 | 1,701.45 | 397.77 | 345.06 | 6,684.27 | 4,959.44 |

**Table 8.** BUSCO analysis of predicted proteins from myxozoan genomes using the eukaryote database. For myxozoans with publicly-available annotations, the predicted protein sets from both the original annotation and from our BRAKER annotation were assessed using BUSCO. N.A. not available. Note 1: We could not find the annotations file from the publication.

| Species | Original annotations |  | BRAKER2 annotations |  |
| --- | --- | --- | --- | --- |
|  | Protein-coding genes | BUSCO (Eukaryota)<br>% where n=225 | Protein-coding genes | BUSCO (Eukaryota)<br>% where n=225 |
| *Sphaeromyxa zaharoni* | N.A. | N.A. | 18,437 | C:35.7 [S:13.7, D:22.0],<br>F:23.5, M:40.8 |
| *Kudoa iwatai* | 5,533 | N.A.<sup>1</sup> | 6,921 | C:67.1 [S:31.0, D:36.1],<br>F:7.5, M:25.4 |
| *Enteromyxum leei* | N.A. | N.A. | 20,185 | C:22.8 [S:21.6, D:1.2],<br>F:24.7, M:52.5 |
| *Myxobolus squamalis* | 5,710 | C:22.0 [S:22.0, D:0.0],<br>F:19.2, M:58.8 | 4,538 | C:29.8 [S:29.8, D:0.0],<br>F:16.9, M:53.3 |
| *Henneguya salminicola* | 8,187 | C:35.3 [S:35.3, D:0.0],<br>F:18.8, M:45.9 | 4,163 | C:32.6 [S:31.0, D:1.6],<br>F:17.3, M:50.1 |
| *Thelohanellus kitauei* | 15,020 | C:29.4 [S:28.6, D:0.8],<br>F:15.7, M:54.9 | 9,498 | C:37.3 [S:34.9, D:2.4 ],<br>F:14.9, M:47.8 |
| *Ceratonova shasta* | N.A. | N.A. | 15,032 | C:27.1 [S:25.1, D:2.0 ],<br>F:8.2, M:64.7 |
| *Myxobolus honghuensis* | 15,433 | C:51.0 [S:49.4, D:1.6],<br>F:14.1, M:34.9 | 16,867 | C:44.4 [S:42.4, D:2.0],<br>F:16.9, M:38.7 |
| *Myxobolus sp.* | - | - | 10,106 | C:44.3 [S:41.6, D:2.7],<br>F:16.5, M:39.2 |

Next, we assigned functional annotations to 57% (5,757 of 10,106) *Myxobolus sp.* proteins with: GO terms to 2,611 proteins; KEGG pathways to 1,873 proteins; and PFAM domains to 4,671 proteins. Nearly all KEGG pathways assigned to at least one *Myxobolus sp.* protein (357/361 KEGG pathways) were common between *Myxobolus sp.* and *M. honghuensis.* Further, 340 pathways contained at least one protein from all myxozoan species. The number of pathways with at least one protein was similar across free-living and parasitic cnidarians, ranging from 349 in *C. shasta* to 382 in *T. kitauei* in myxozoans and from 390 in *C. hemisphaerica* to 405 in *S. pistillata.* When we looked at the 339 KEGG pathways with at least one protein from all myxozoan and free-living cnidarian species, we observed that, on average, free-living cnidarians had more proteins assigned to the pathway compared to myxozoans.

We assigned 1,729 PFAM domains to at least one *Myxobolus sp.* protein, of which 1,522 domains (88%) were also found in the genome of *M. honghuensis.* Of these 1,729 domains, nine were not identified in any of the free-living cnidarian species (Table 9). Ten proteins had the “Arg_repressor” (Arginine repressor, DNA binding domain) protein domain in *Myxobolus sp.* and *C. shasta,* two in *T. kitauei* and none in any free-living cnidarians analyzed in this study.

**Table 9.** Number of myxozoan proteins containing the nine domains that were identified in at least one *Myxobolus sp.* protein but none of the free-living cnidarians used in the study

| Pfam domain identifier | *Myxobolus sp.* | *M. honghuensis* | *M. squamalis* | *C. shasta* | *E. lei* | *H. salmnicola* | *K. iwatai* | *S. zaharoni* | *T. kitauei* |
| --- | --- | --- | --- | --- | --- | --- | --- | --- | --- |
| Arg_repressor | 10 | 0 | 0 | 10 | 0 | 0 | 0 | 0 | 2 |
| Terminase_5 | 4 | 0 | 0 | 0 | 0 | 0 | 0 | 0 | 0 |
| DUF1398 | 1 | 0 | 0 | 0 | 0 | 0 | 0 | 0 | 0 |
| Lambda_Bor | 1 | 0 | 0 | 0 | 0 | 0 | 0 | 0 | 0 |
| Mononeg_RNA_pol | 1 | 0 | 0 | 0 | 0 | 0 | 0 | 3 | 0 |
| Phage_holin_3_1 | 1 | 0 | 0 | 0 | 0 | 0 | 0 | 0 | 0 |
| Phage_lysis | 1 | 0 | 0 | 0 | 0 | 0 | 0 | 0 | 0 |
| RdRP_2 | 1 | 0 | 0 | 0 | 0 | 0 | 0 | 0 | 0 |
| Saw1 | 1 | 0 | 0 | 0 | 0 | 0 | 0 | 0 | 0 |

A further 22 PFAM domains were present in more *Myxobolus sp.* proteins compared to free-living cnidarians (the number of proteins with a domain in *Myxobolus sp.* was at least three times the average number of proteins containing the domain in free-living cnidarian species). From these 22 domains, “RVP_2”, “DDE_Tnp_IS1595”, and “FLYWCH” were the top three most abundant protein domains. Proteins with the FLYWCH domain are a part of a superfamily of DNA transposon-derived conserved families of transcription factors. The number of FLYWCH domains ranged from none in three myxozoans to 55 in *Myxobolus sp.* in myxozoans, and none in two free-living cnidarians to three in *S. pistillata* in free-living cnidarians. In *Myxobolus sp.,* the majority of proteins with the FLYWHC domain also contained the MULE transposase protein domain.

We found that both *Myxobolus sp.* and *M. honghuensis* had more proteins containing the “Hexokinase_1” and “Hexokinase_2” protein domains when compared to free-living cnidarians. *Myxobolus sp.* had six proteins containing both domains. Phylogenetic analysis of all proteins that contained the two domains showed that the family underwent duplication in these two species (Figure 2).

**Figure 2.**
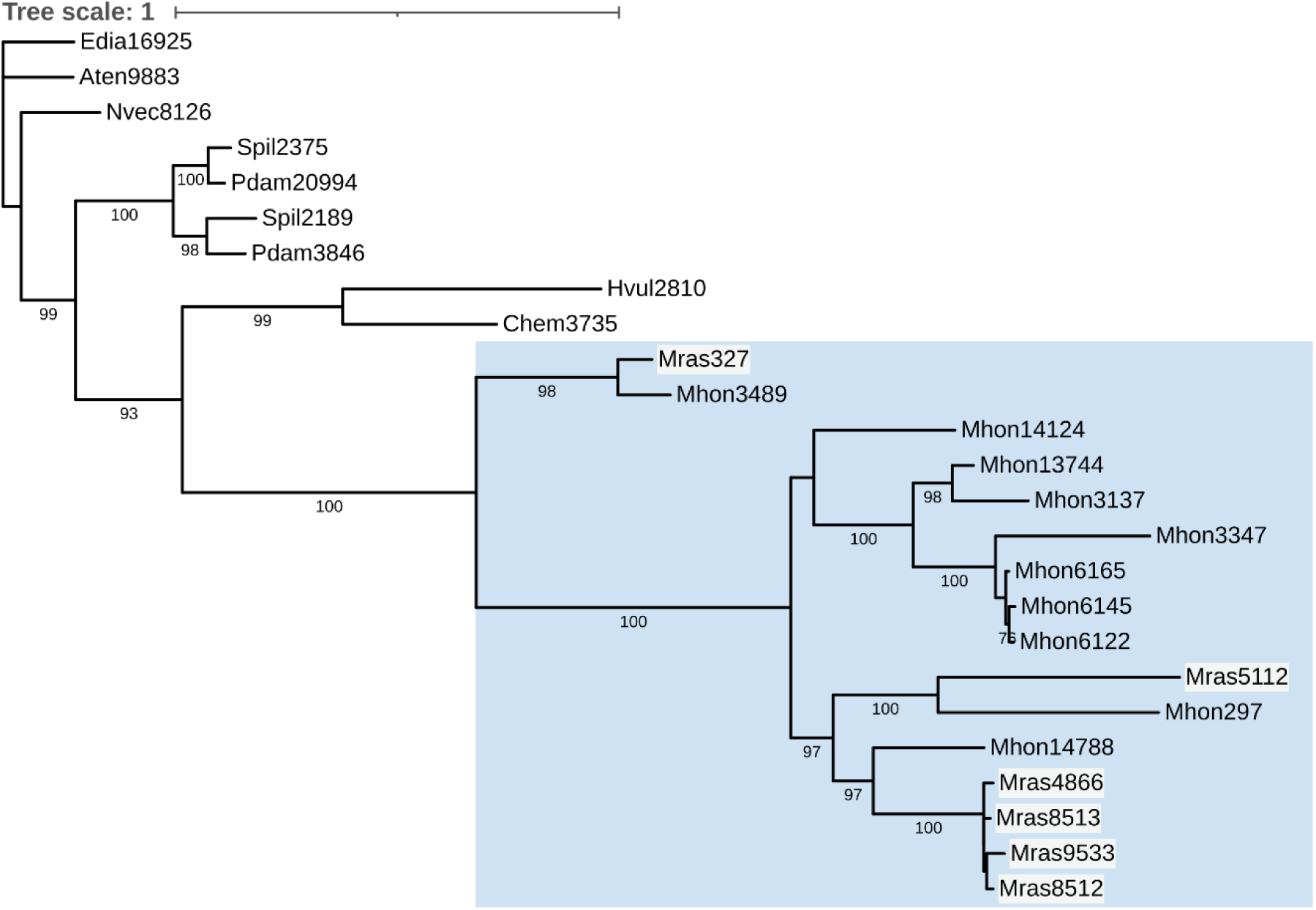
Phylogenetic tree of proteins containing the “Hexokinase_1” and “Hexokinase_2” protein domains in free living cnidarians and *Myxobolus* sp. and *M. honghuensis*, in which a blue box represents the clade containing all myxozoan proteins and *Myxobolus* sp. proteins have a white background. Bootstrap support values of 75% and above are annotated on the branches of the tree.

### 3.3. Gene family analysis

Using OrthoFinder, we identified 9,843 groups of orthologous proteins (OGs) that contained at least one myxozoan protein, of which 3,972 OGs contained at least one *Myxobolus sp.* protein. Of these, 3,095 OGs also contained protein from *M. honghuensis*. There were 877 OGs with at least one *Myxobolus sp.* protein but not *M. honghuensis,* and 1070 OGs vice versa. Less than one-third of all 9,843 OGs (2,727 OGs) contained a protein from the majority of myxozoan species used in the analysis (four of the seven). These 2,727 OGs represent a set of core conserved proteins in myxozoans.

#### 3.3.1. Phylogenetic relationships between the myxozoan species

Using the 57 single-copy orthologs, we reconstructed a ML-based phylogeny of both myxozoans and free-living cnidarians and found that *M. honghuensis* is the sister species to *Myxobolus sp.* with the other relationships agreeing with previous studies (Figure 3).

**Figure 3.**
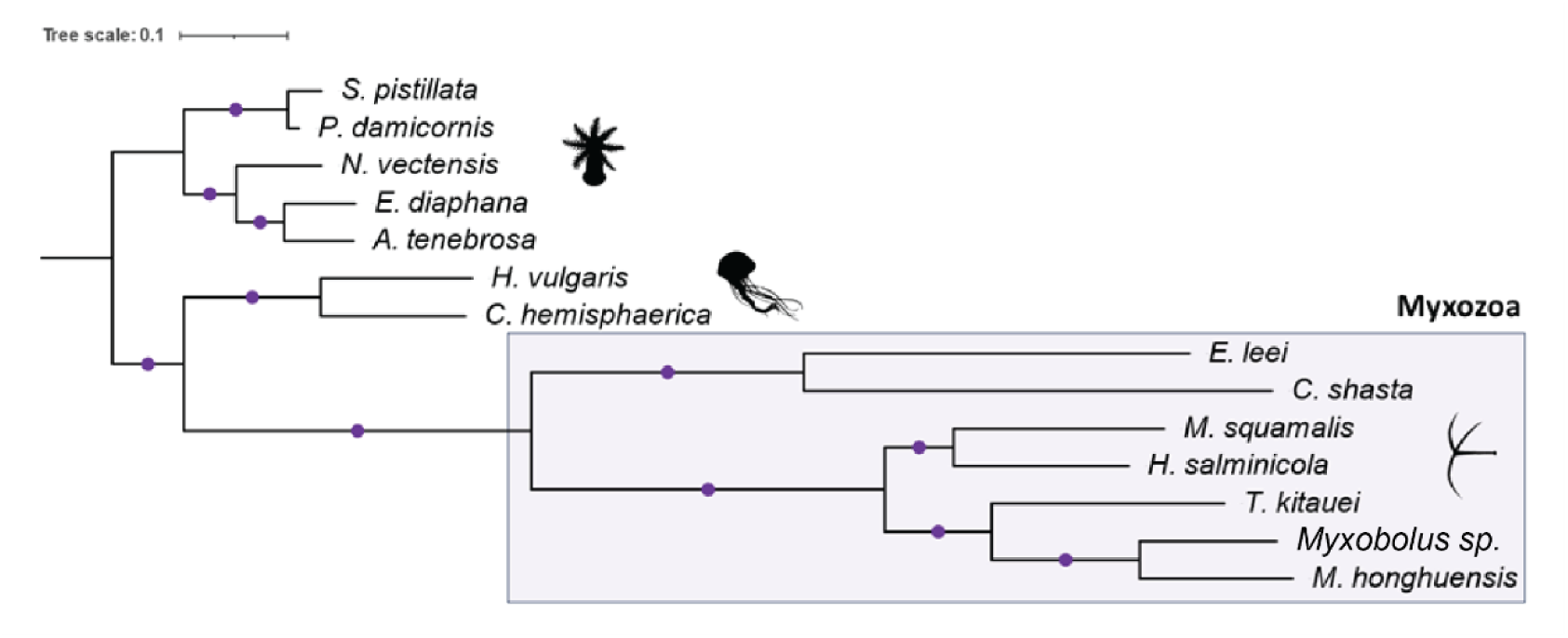
Phylogenetic relationships between free-living and parasitic cnidarians generated using IQTREE. Single-copy orthologs identified by OrthoFinder were used as input. All nodes with boot-strap values greater than 90% are annotated with purple circles. The silhouettes for Anthozoa, Medusozoa, and Myxozoa are taken from PhyloPic (www.phylopic.org).

#### 3.3.2. Gene family expansions in *Myxobolus sp*

Using the species tree and full set of orthology clusters results, we identified 10 gene families that were significantly expanded in myxozoans. We identified a further 19 gene families that were significantly expanded in only *Myxobolus sp.*, the majority of which were involved in repeat-related processes, and hence were not analyzed further. Here, we describe on three gene families of interest.

The first gene family of interest (OG0000160) was comprised of the orthologs of the *H. vulgaris* SERPINB6 (serine protease inhibitor B6). This family underwent expansion on the branch leading to *Myxobolus sp., M. honghuensis,* and *T. kitauei,* which was followed by a contraction on the branch leading to *M. honghuensis* (Figure 4A). All myxozoan proteins in this OG clustered together in the maximum likelihood (ML) phylogenetic tree (Figure 5). Since serpins are considered as potential targets for myxozoan control and have been extensively studied in *M. cerebralis*, we inspected this gene family further. We observed that only 11 of the 25 *Myxobolus sp.* proteins from this OG contained the “Serpin” domain. Additionally, two more OGs contained *Myxobolus sp.* proteins with the “Serpin” domain. Of these, one OG (OG0022393) had proteins only from *Myxobolus sp.* and *M. honghuensis.* We analyzed the phylogenetic relationships between all proteins containing serpin domains from *Myxobolus sp.*, *M. honghuensis, M. cerebralis,* and free-living cnidarians. We observed that all the myxozoan serpins clustered together (Figure 6). Further, myxozoan homologs of *M. cerebralis* serpin 1 underwent expansion in *Myxobolus sp.* and *M. honghuensis*.

**Figure 4.**
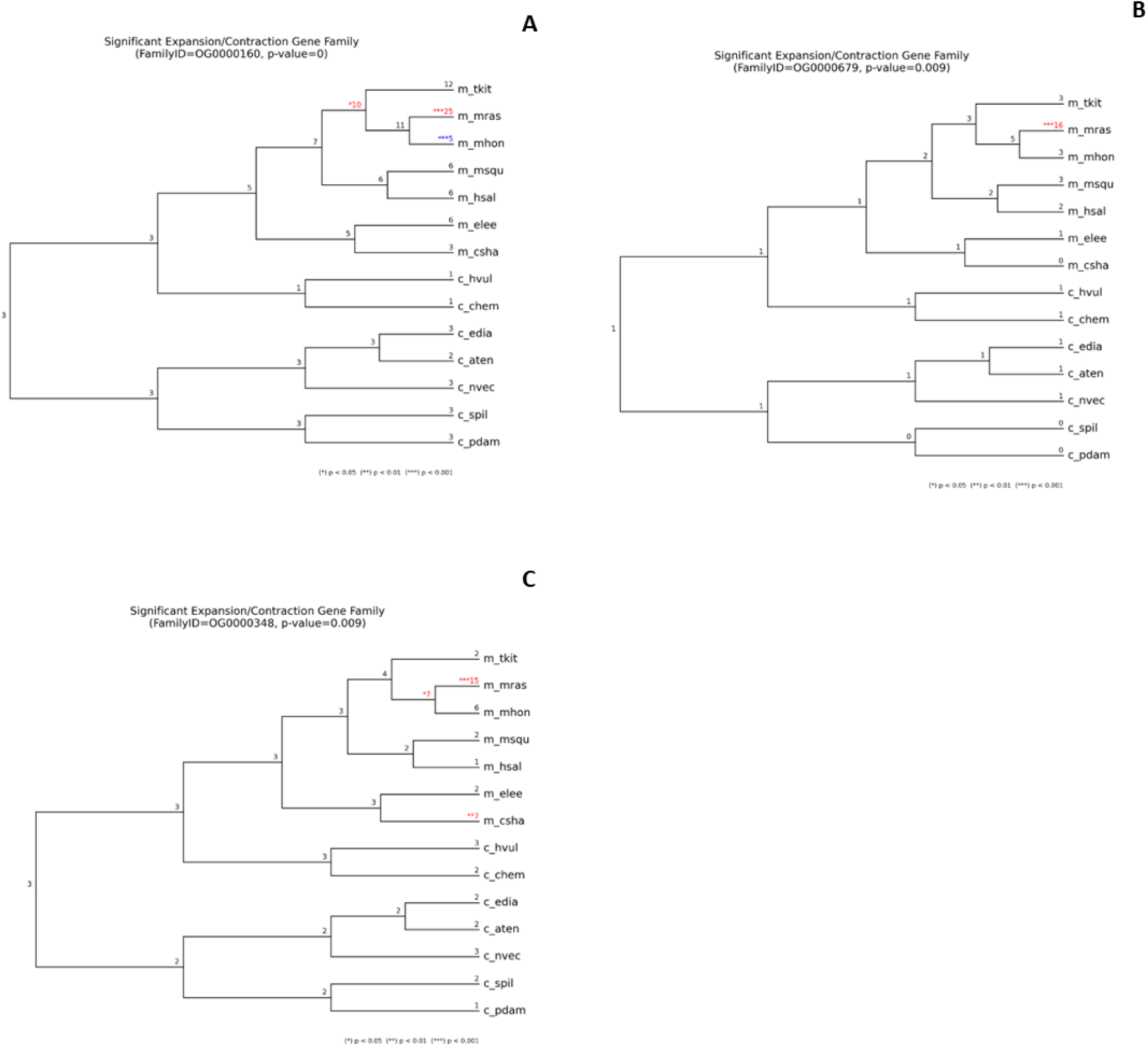
CAFE5 analysis of A) OG0000160, B) OG0000679, and C) OG0000348 with significant expansions and contractions colored in red and blue, respectively. Significant changes on the branch are labelled with asterisks, where red color indicates gene family expansion and blue color indicates gene family contraction. (*) indicates a p-value < 0.05, (**) indicates a p-value < 0.01, and (***) indicates a p-value < 0.001.

**Figure 5.**
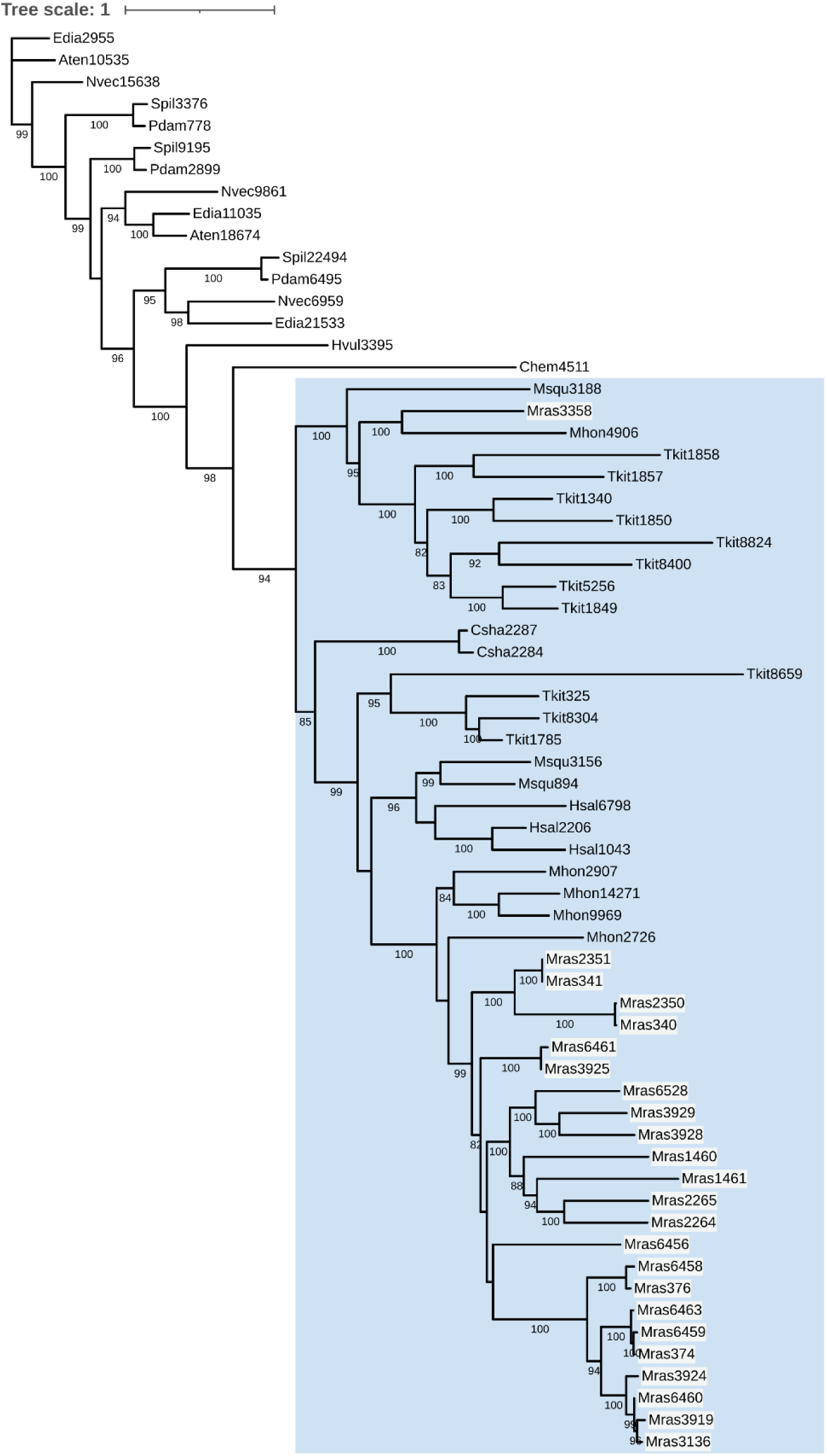
Phylogenetic tree of the gene family OG000160, in which the *Myxobolus sp.* proteins have a white background and a blue box represents the clade containing all myxozoan proteins. Bootstrap support values of 75% and above are annotated on the branches of the tree.

**Figure 6.**
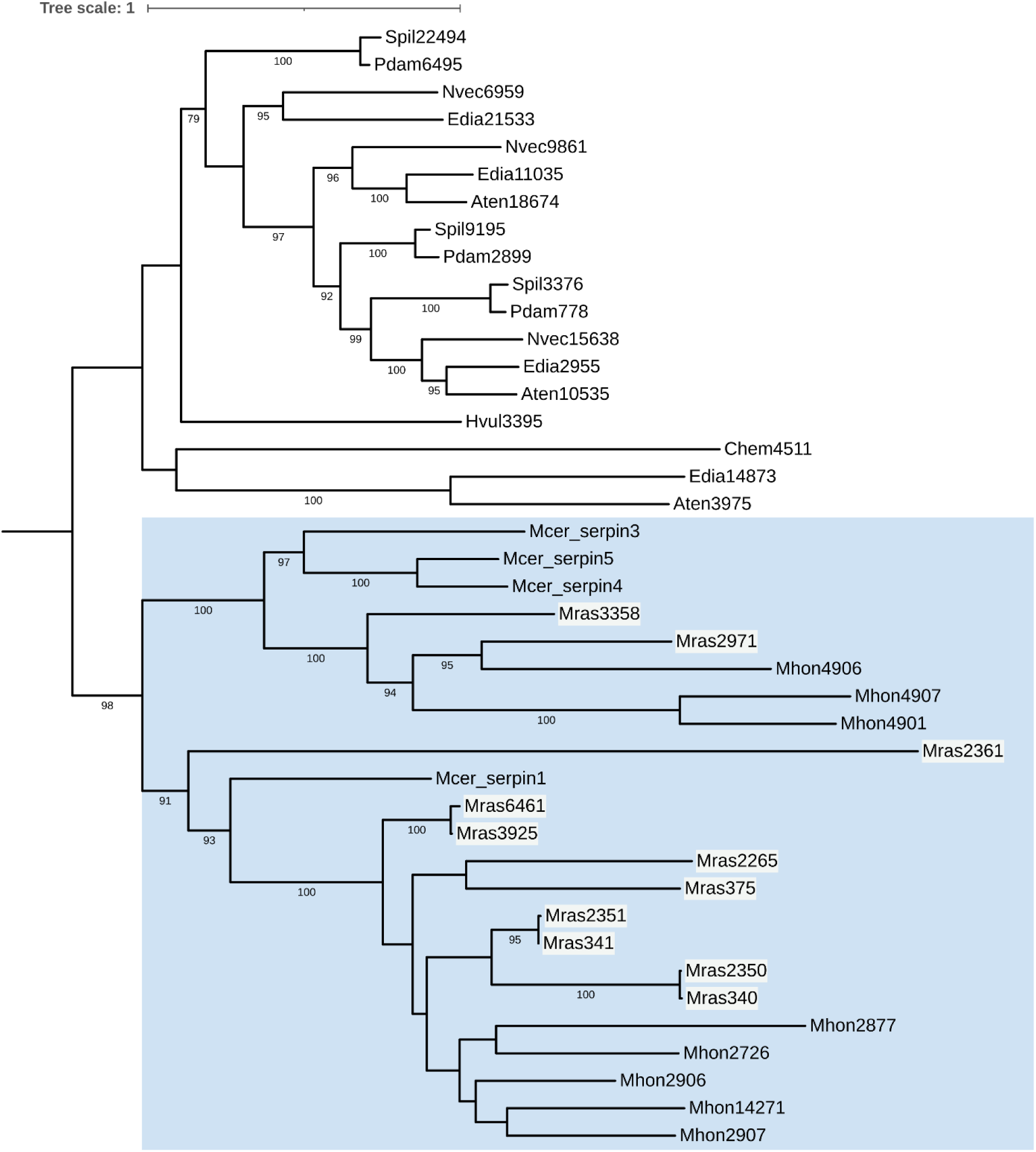
Phylogenetic tree of proteins containing the “Serpin” domain in free-living cnidarians, *Myxobolus sp.*, *M. honghuensis*, and *M. cerebralis* in which the *Myxobolus sp.* proteins have a white background and a blue box represents the clade containing all myxozoan proteins. Bootstrap support values of 75% and above are annotated on the branches of the tree.

The second gene family of interest (OG0000679) underwent expansion in *Myxobolus sp.* and was comprised of orthologs of the *H. vulgaris* protein “Uncharacterized protein LOC100211173 isoform X1”. Proteins in this OG harbor the “Lupus La” protein domain. This gene family experienced an expansion on the branch leading to both *Myxobolus sp.* and *M. honghuensis* (Figure 4B, 7). Finally, the third gene family of interest that experienced expansions in *Myxobolus sp.* consisted of orthologs of three peroxiredoxin proteins in *H. vulgaris*. This gene family experienced an expansion on the branch leading to *Myxobolus sp., M. honghuensis,* and *T. kitauei,* and also in *C. shasta* (Figure 4C, 8).

**Figure 7.**
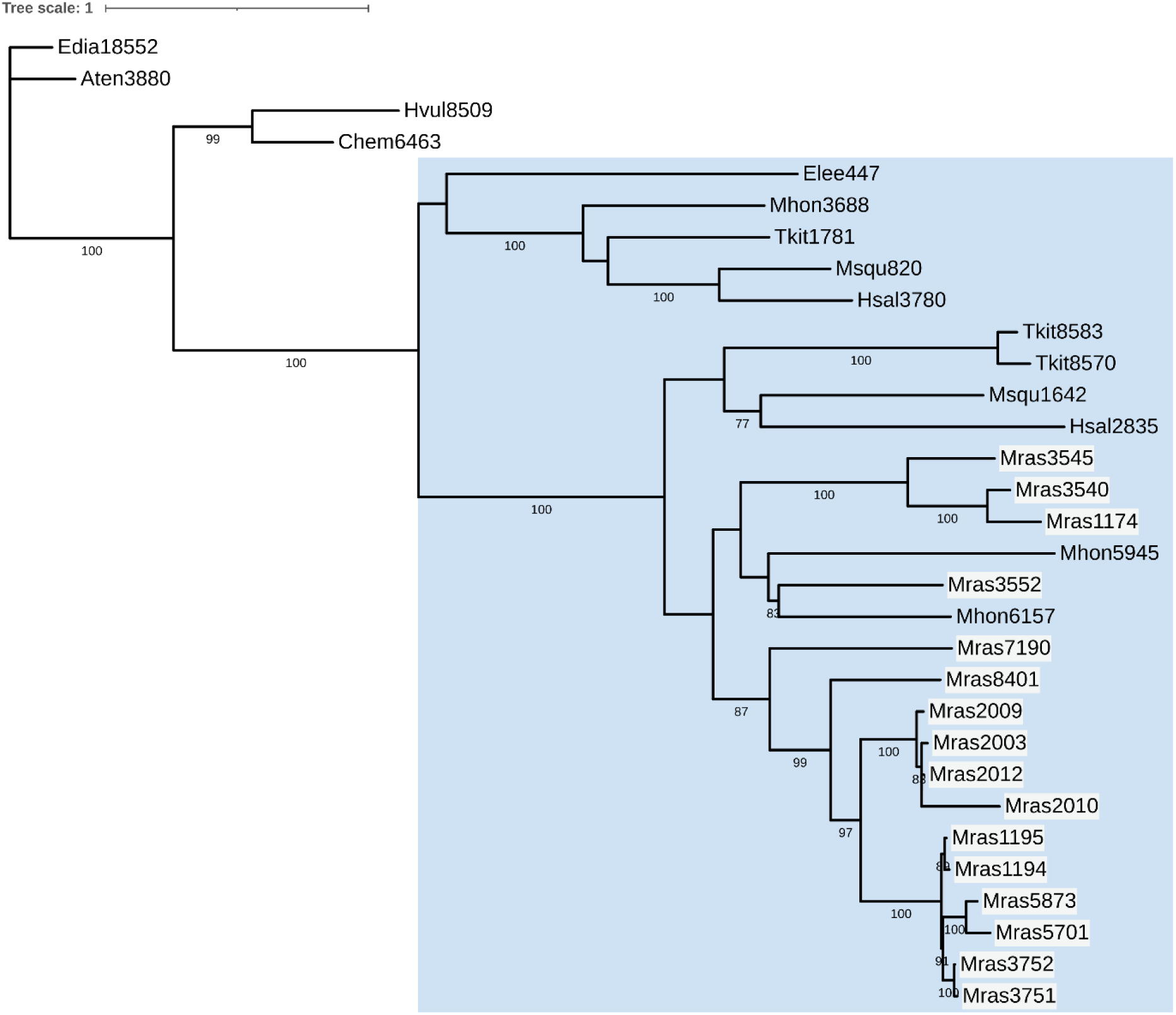
Phylogenetic tree of the gene family OG000679 in which the *Myxobolus sp.* proteins have a white background and a blue box represents the clade containing all myxozoan proteins. Bootstrap support values of 75% and above are annotated on the branches of the tree.

**Figure 8.**
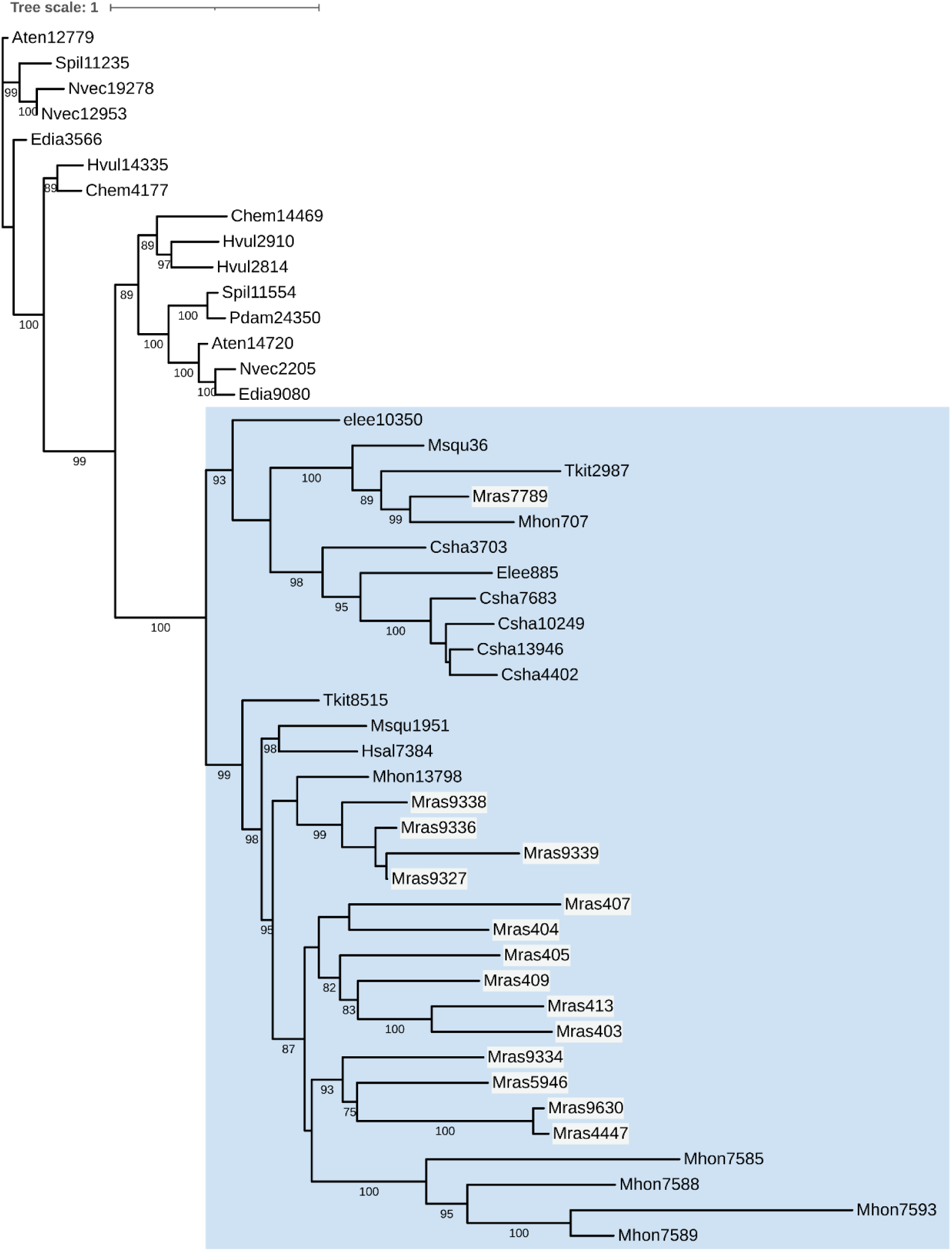
Phylogenetic tree of the gene family OG000348 (peroxiredoxin) in which the *Myxobolus sp.* proteins have a white background and a blue box represents the clade containing all myxozoan proteins. Bootstrap support values of 75% and above are annotated on the branches of the tree.

#### 3.3.3. Gene families of interest in *Myxobolus sp*

In addition to these three expanded gene families, we explored the evolution of two gene families that have been of interest in myxozoan biology. First, we analyzed the complement of proteases found in *Myxobolus sp.* since proteases have been proposed to be promising targets for myxozoan control. We observed that 377 *Myxobolus sp.* proteins had a top hit to 155 proteases from the MEROPS database: 39 metalloproteases, 38 cysteine proteases, 22 serine proteases, 19 threonine proteases, 17 aspartic proteases, and 20 protease inhibitors. The most abundant category of proteases was A28B (the aspartic peptidase “skin SASPase”), followed by S01A (serine peptidase “chymotrypsin A”), and C01A (cysteine peptidase “papain”). We also identified 31 proteins from the I04 family of serine and cysteine endopeptidase inhibitors.

We aligned all but 16 of these 377 *Myxobolus sp.* proteases to proteins from the fathead minnow and 13 of the remaining 16 proteins could be aligned to Actinopterygii (ray-finned fish) proteins from the NR database (e-value ≤ 1e^-3^). We inspected the function annotation remaining three proteins (mras5248, mras6483, and mras9733): protein mras5248 belonged to the same homologous superfamily as the mouse retroviral-like aspartic protease 1 (A28B subfamily) and also has the same PFAM protein domain (“gag-asp_protease” gag-polyprotein putative aspartyl protease); protein mras6483 was assigned to the same homologous superfamily as the mouse retroviral-like aspartic protease 1 (A28B subfamily) but did not have any predicted PFAM protein domains; protein mras9733 could not be assigned any family membership or PFAM domains.

Next, we analyzed the repertoire of minicollagen and nematogalectin proteins in *Myxobolus sp.*. We searched for significant sequence similarity with minicollagen sequences from myxozoans and free-living cnidarians, and analyzed phylogenetic relationships between them (Figure 9).

**Figure 9.**
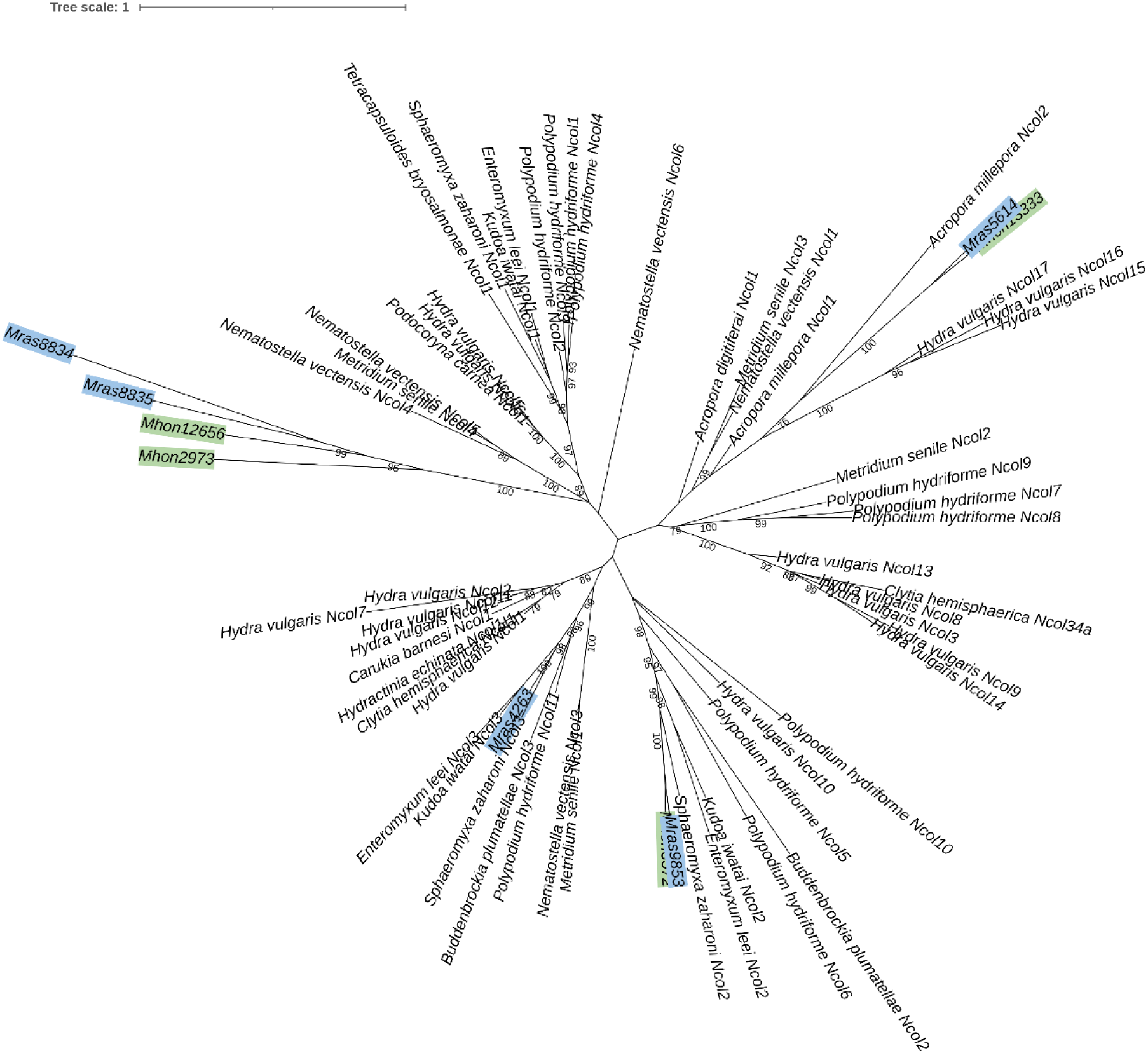
Phylogenetic tree of the minicollagens, in which *Myxobolus sp.* and *M. honghuensis* sequences are labelled with blue and green color respectively. Bootstrap support values of 75% and above are annotated on the branches of the tree.

Previously, minicollagens were shown to belong to three different clades: I, II, and III. We found one *Myxobolus sp.* protein in each of clades II and III, and none from clade I. In addition, we identified one *Myxobolus sp.* and one *M. honghuensis* protein that grouped with free-living cnidarian minicollagens from clade III. There were two *Myxobolus sp.* proteins and two *M. honghuensis* proteins that clustered together but did not have sufficient phylogenetic support to assign to a minicollagen clade. For the nematogalectins, we found three proteins in *Myxobolus sp.*: one closely related to *M. bejeranoi* nematogalectin A; one to *M. bejeranoi* nematogalectin C; and the third to a putative nematogalectin in *S. zaharoni* (Figure 10).

**Figure 10.**
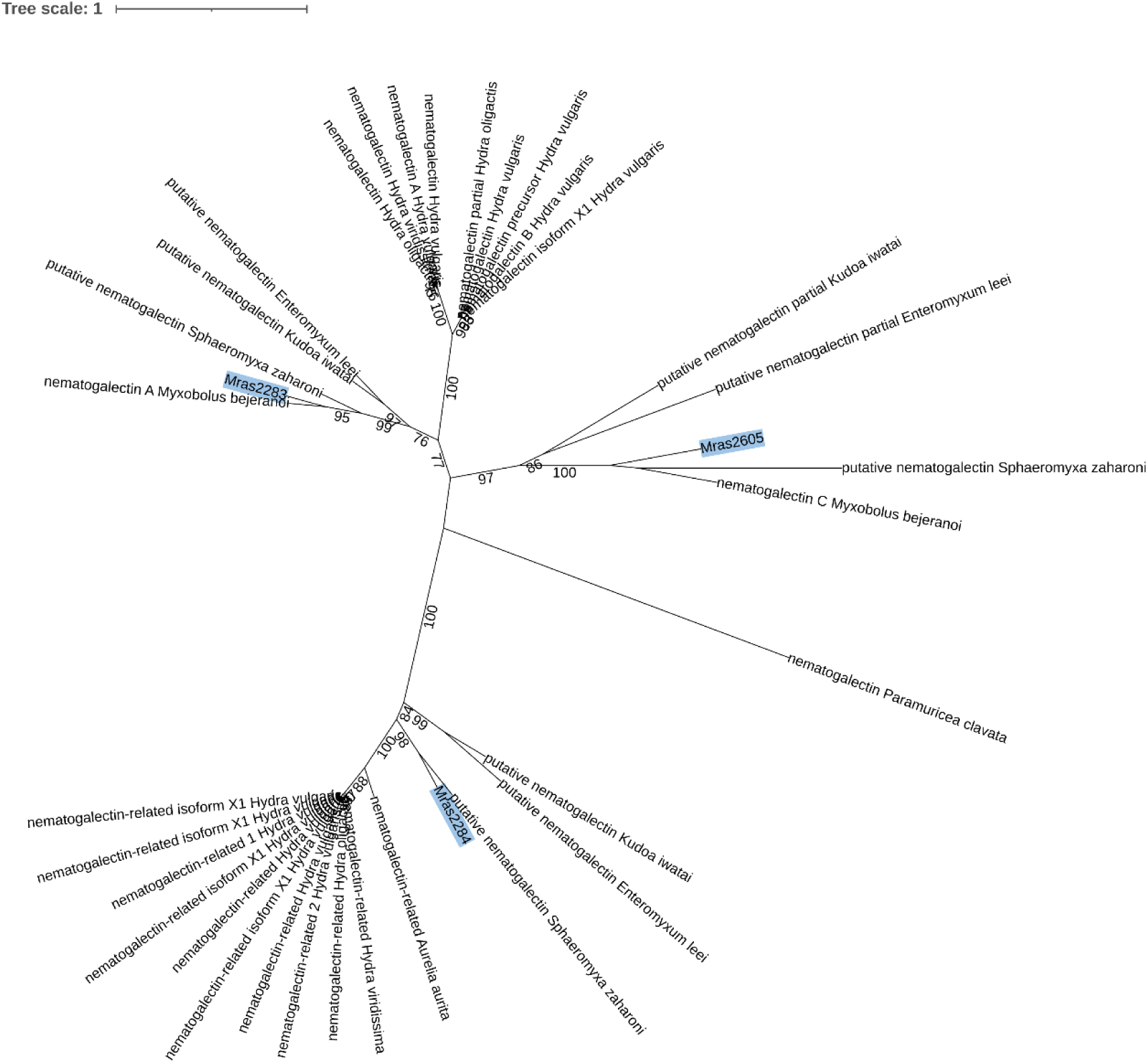
Phylogenetic tree of the nematogalectins in which *Myxobolus sp.* sequences are labelled with blue. Bootstrap support values of 75% and above are annotated on the branches of the tree.

#### 3.3.4. Gene-loss in *Myxobolus sp*

We identified 111 OGs with proteins from *M. honghuensis* and the majority of the remaining five myxozoan species, but none from *Myxobolus sp..* We extracted the 178 *M. honghuensis* proteins from these 111 OGs and examined the protein domains and KEGG pathways associated with these proteins as determined by EggNOG-mapper. The top three most abundant protein domains were “Dimer_Tnp_hAT” (hAT family C-terminal dimerisation region), “Pkinase” (Protein kinase domain), and “UAA” (UAA transporter family). The top three most abundant KEGG pathways were “Metabolic pathways” (map01100), “Cellular senescence” (map04218), and “cGMP-PKG signaling pathway” (map04022).

#### 3.3.5. *Myxobolus sp.* proteins with no sequence similarity to proteins from the fathead minnow

We detected 3,444 *Myxobolus sp.* proteins that did not have sequence similarity to any fathead minnow protein, as identified using BLASTP (e-value ≤ 1e^-3^), of which 639 proteins also had BLASTP hits to proteins from Actinopterygii species (e-value ≤ 1e^-3^). The remaining 2,805 *Myxobolus sp.* proteins were queried against the ChEMBL database to search for potential drug targets. To this end, we identified eight *Myxobolus sp.* proteins which had a BLASTP hit to a protein from the ChEMBL database (e-value ≤ 1e^-5^). For a more stringent selection criteria, we only looked at *Myxobolus sp.* proteins that did not have a BLASTP hit to proteins from the fathead minnow (e-value ≤ 0.05). This yielded three proteins which we analyzed in detail.: “mras18” was aligned to inositol phosphorylceramide synthase (CHEMBL1641343), “mras2094” to gag-pol polyprotein (CHEMBL3638331), and “mras2163” to heat shock protein hsp-16.2 (CHEMBL2146313).

### 3.4. Characterization of the mitochondrial genome and mitochondrial proteome

We identified a single 31,057 bp contig as the mitochondrial genome (mt-genome) of *Myxobolus sp.* (Figure 11A). The *Myxobolus sp.* mt-genome was predicted to encode only five of the 13 protein-coding genes that are typically found in most animal mitochondrial genomes—*cox1*, *cox2*, *nad1*, *nad5*, and *cob*—as well as two ribosomal RNA genes (*rrnS* and *rrnL*). The *rrnL* gene was split into two fragments. We aligned both the fragments of the 16S RNA gene (*rrnL*) to the NT database and found that the top hit to both fragments was from the myxozoan *Myxobolus squamalis*. When we searched the nuclear genome of *Myxobolus sp.* with the same HMM profiles of protein-coding genes encoded in the mt-genome, we found that *atp6* was transferred to the nuclear genome and had acquired introns and an mitochondria-targeting signal (MTS). In fact, we found that *atp6* was transferred to the nuclear genome and acquired an MTS in *M. honghuensis*, *S. zaharoni*, *T. kitauei, K. iwatai*, and *M. squamalis*. None of the other protein-coding genes in the mt-genome were transferred to the nuclear genome in myxozoans.

**Figure 11.**
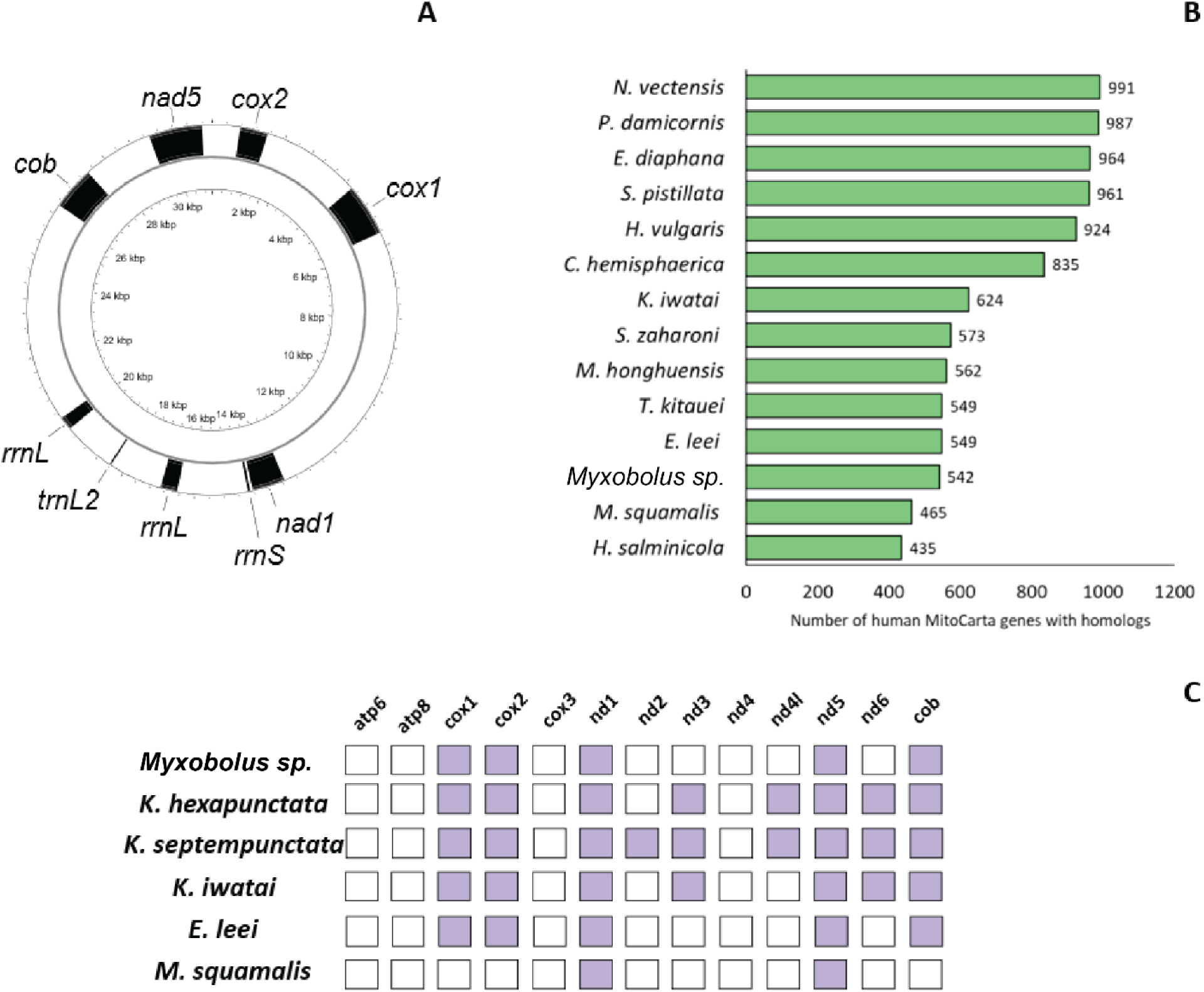
A) The mitochondrial genome of *Myxobolus sp.* visualized using PROKSEE, B) The number of human MitoCarta genes with Blast hits to myxozoan (purple) genomes and cnidarian (green) genomes, C) The presence/absence of protein-coding genes in the mitochondrial genomes of myxozoans.

Next, we analyzed the mitochondrial proteome—the set of proteins encoded in the nuclear genome and imported into the mitochondrion—of *Myxobolus sp.*. First, we identified homologs of nuclear-encoded mitochondrial proteins from humans in the cnidarian genomes. For this, we aligned the set of human MitoCarta v3 proteins to the proteomes from myxozoans and free-living cnidarians. *Myxobolus sp.* had homologs of 542 of the 1,136 human MitoCarta genes (Figure 11B). On average, myxozoans had homologs of 526 MitoCarta genes, compared to 951 for the free-living cnidarians. We identified 741 MitoCarta genes that had homologs in at least one myxozoan species, of which 241 were present in all. We used ConsensusPathDB to find out the functions of these 241 mitochondrial protein-coding genes. The top five enriched KEGG pathways included “Valine, leucine and isoleucine degradation”, “TCA cycle”, “Fatty acid degradation”, and “Aminoacyl-tRNA biosynthesis”. On the contrary, 395 mitochondrial protein-coding genes did not have a significant sequence similarity hit to any myxozoan protein in this study, representing a potential loss of mitochondrial functions in myxozoans. ConsensusPathDB found that the top five enriched pathways in this missing set included “Oxidative phosphorylation”, “Thermogenesis”, and “Apoptosis”.

Next, we identified potential mitochondria-targeted proteins in *Myxobolus sp.* which do not have homologs to known mitochondrial proteins in humans. For this, we identified *Myxobolus sp.* proteins that are predicted to possess a mitochondrial targeting signal (MTS) at their N-terminus end using MitoFates. We observed that only 271 *Myxobolus sp.* proteins were predicted to harbor an MTS, of which 36 were predicted to possess an MTS with a high probability (at least 90%).

Of these, 15 had a hit to a human protein, for which the top human hit was also predicted to possess a MTS. A further seven *Myxobolus sp.* proteins had a hit to a human protein that did not have MTS. These proteins represent potential instances of neolocalization in the parasite (an instance where the homolog of a non-mitochondrial protein in human is targeted to the mitochondria in *Myxobolus sp.*). These included the endogenous retrovirus group K member 113 Pol protein (P63132), NYNRIN (Q9P2P1), alpha-1,3-mannosyl-glycoprotein 2-beta-N-acetylglucosaminyltransferase (P26572), adenosine 5’-monophosphoramidase HINT1 (P49773), and cytoplasmic asparagine--tRNA ligase (O43776). The remaining 14/36 proteins did not have BLASTP hit to any human protein, and we used BLASTP to search for proteins with sequence similarity in the NR database. Of these, five did not have a BLASTP hit to any non-myxozoan protein in the NR database.

## 4. Discussion

### 4.1. The high repeat content of the *Myxobolus* sp. genome contributes to genome expansion and necessitates the use of long-read sequencing of myxozoan genomes

In this study, we sequenced, annotated, and analyzed the genome of *Myxobolus* sp., a recently discovered myxozoan parasite of fathead minnows in southern Alberta. We used the Oxford Nanopore long-read sequencing technology, and thus were able to provide the second long-read genome in this species-rich group of parasites. While relatively short for an animal genome, it is one of the most repetitive. The only other long-read-based myxozoan genome, *M. honghuensis*, is also highly repetitive. Such repetitive sequences can negatively affect the assembly of a genome, especially in genomes sequenced using short-read platforms where repetitive loci are more likely to be misassembled, which likely explains the relatively high number of scaffolds in both the *M. honghuensis* and our *Myxobolus sp.* assemblies (Tørresen et al., 2019). Since both long-read genomes belong to a single genus in myxozoans, we cannot say with confidence whether high repeat content is a feature of all myxozoan genomes.

The high repeat content in the *Myxobolus sp.* genome is a significant factor explaining the large genome size in this species. We detected a proliferation of the MULE-MuDR elements in the genome. A recent analysis of the diversity of these Mutator-like elements (MULEs) across eukaryotes found that, outside of plants, many genomes containing these elements were from pathogens (Dupeyron et al., 2019). Indeed, in fungal pathogens, such TE-expansions have been proposed to account for genome expansion (Raffaele & Kamoun, 2012). Furthermore, the increase in repeat content in parasites could result in new functions and genome arrangements which benefit the parasite’s efforts to escape host defenses (Dong et al., 2015). It is possible that the small sizes of myxozoan genomes that were sequenced using short-read sequencing platforms, and the smaller repeat content predicted in them, are an underestimation of the true genome size of these parasites.

The increase in repeat content of the *Myxobolus sp.* genome can also further our understanding of myxozoan biology. A good example of this is the expansion in proteins with the FLYWCH domains identified in *Myxobolus sp.*. A recent analysis of DNA transposon-derived transcription factors in animals found that FLYWCH-domain containing transcription factors are rare in animals but have undergone expansion in one phylum of non-bilaterian animals – Ctenophora (Mukherjee & Moroz, 2023). This study did not include myxozoans in their comparisons. We show that *Myxobolus sp.* could contain the highest repertoire of these FLYWCH proteins in animals.

### 4.2. Gene family expansions

A prominent feature of myxozoan genome evolution is the loss of genes that are otherwise well-conserved in free-living eukaryotes. Hence, gene families that defy this trend and instead undergo expansion are interesting as they represent potential adaptations to the parasitic myxozoan lifestyle.

In one such example of gene family expansion, serpins (serine proteases) play an important role in parasitic organisms, like defense against host proteases in *Schistosoma mansoni* and acting as immunomodulators and inhibit host immune responses in nematodes (Maizels et al., 2001; Valdivieso et al., 2015). Recently, a study analyzed the expression-profiles of *M. cerebralis* serpins and proposed that they represent good targets for inactivating the parasite at an early stage in the fish host (Eszterbauer et al., 2021). We found a group of *Myxobolus sp.* proteins that clustered together with serpins 3, 4, and 5 from *M. cerebralis.* Researchers have shown that the expression of these three serpins is upregulated during fish invasion and early development, and their functional predictions suggest that these three are caspase inhibitors. Thus, the *Myxobolus sp.* proteins that cluster together with these three serpins represent promising targets for parasite control. Additionally, serpin 1, predicted to be a putative chymotrypsin-like inhibitor, has undergone extensive duplication in *Myxobolus sp..* While this suggests that the homologs of serpin 1 are probably involved in important functions in *Myxobolus sp.,* developing targets against them would be challenging.

Further, we also identified an expansion in a family of peroxiredoxins in *Myxobolus sp.* and other myxozoans. Peroxiredoxins are ubiquitous enzymes with roles in antioxidation and signaling systems (Gretes et al., 2012). These enzymes play important roles in parasitic defenses against reactive oxygen and nitrogen species generated by the host immune system. These enzymes have been proposed to be drug targets several parasites like trypanosomes and as vaccine antigens in *L. major* (Bayih et al., 2014).

We also observe that both *Myxobolus sp.* and *M. honghuensis* have a higher number of hexokinase domain containing proteins. Hexokinases catalyze the first step of glycolysis. The increase in the hexokinase repertoire in the parasite could reflect the reliance of the parasite on glycolysis to compensate for a reduced and inefficient oxidative phosphorylation system. Indeed, we noticed that several human OXPHOS proteins did not have homologs in *Myxobolus sp.*. These could represent a lead for future antiparasitic strategy.

### 4.3. *Myxobolus* sp. has a reduced mitochondrial genome and proteome

Most animal mitochondrial genomes are single circular compact molecules that encode 37 genes (13 protein coding genes, 2 rRNA genes, and 22 tRNA genes) (Lavrov & Pett, 2016). Myxozoan mitochondrial genomes differ from this archetypal animal mitochondrial genome in several aspects. For instance, all myxozoan genomes have a reduced set of genes (reviewed in (Lavrov & Pett, 2016)). *Enteromyxum leei* has a multipartite mitochondrial genome, comprised of eight megacircles (Yahalomi et al., 2017). The mitochondrial genome of *Myxobolus sp.* is no exception. It represents just the second mitochondrial genome from the species-rich *Myxobolus* genus. The only other mitochondrial genome from the genus belongs to *M. squamalis,* and only a single protein coding gene can be identified in the genome. Further, while researchers produced a high-quality nuclear genome of *M. honghuensis*, they were unable to identify a contig containing the mitochondrial genome. Thus, here we present the most complete mitochondrial genome from the genus *Myxobolus*. It is worth noting that while the mitochondrial genome of *Myxobolus sp.* lacks several protein-coding genes, it is larger than the human mitochondrial genome, due to large intergenic regions, instead of one large non-coding region in most animal mitochondrial genomes. Further, myxozoan mitochondrial genomes have some of the fastest rates of sequence evolution in animals (Lavrov & Pett, 2016), which could make identifying some mitochondrial genes difficult. In fact, some myxozoan mitochondrial genomes from the genus *Kudoa* have some unidentified open reading frames (ORFs) (Takeuchi et al., 2015), which could be standard animal mitochondrial genes that have evolved beyond recognition by sequence-based approaches.

We find that the mitochondrial gene *atp6* is transferred to the nuclear genome in multiple myxozoans and has acquired a mitochondrial targeting signal. This suggests that the protein encoded by this gene is imported into the organelle, however proteomic analyses need to be done to confirm this. Such a transfer of *atp6* to the nuclear genome and the acquisition of a mitochondrial targeting signal has been reported before, like in the ctenophore *Mnemiopsis leidyi* (Pett et al., 2011) and in the non-animal eukaryote *Chlamydomonas reinhardtii* (Funes et al., 2002). This transfer of *atp6* to the nuclear genome and its potential for its subsequent import back into the organelle make myxozoans potential models to study mitochondrial protein import for aiding efforts in mitochondrial gene therapy (Figueroa-Martínez et al., 2011; Manfredi et al., 2002), as was suggested for *M. leidyi* (Pett et al., 2011).

All proteins encoded by genes in the mitochondrial genome function in complexes that require proteins that are encoded in the nuclear genome and imported into the organelle. The majority of mitochondrial processes rely on just the import of nuclear-encoded proteins. This underscores the need to analyze the mitochondrial proteome for analyzing the evolution of mitochondrial processes in myxozoans. The analysis of mitochondrial proteomes outside of model animals is limited. Only a few studies have looked at the evolution of the mitochondrial proteome in myxozoan species, both showing that myxozoans lack several mitochondrial proteins found in mammals (Muthye & Lavrov, 2018; Yahalomi et al., 2020). Our analysis of the mitochondria proteomes from myxozoans align with these results. On average, just less than half the mitochondrial proteins from mammals had homologs in myxozoans, compared to ∼85% in the three non-parasitic cnidarians. We found that around 35% of the human mitochondrial proteins were missing in all the myxozoan species analyzed, representing a set of core lost mitochondrial functions. These missing proteins were enriched in pathways such as oxidative phosphorylation and apoptosis. These findings are consistent with a recent study that found that several apoptosis proteins are lost in myxozoan genomes (Neverov et al., 2023).

### 4.4. *Myxobolus* sp. represents a potential model system to study host-parasite interactions in the genus *Myxobolus*

The lack of experimentally confirmed life cycles from myxozoans impedes efforts to analyze how these parasites infect the hosts, transmit between them, and interact with their hosts post infection. Within the Myxozoa, only six life cycles have been established in the laboratory, of which three are from the genus *Myxobolus*: *M. cerebralis* (El-Matbouli et al., 1999; Wolf & Markiw, 1984), *Myxobolus pseudodispar* (Kallert et al., 2007; Marton & Eszterbauer, 2012; Székely et al., 1999, 2001), and *Myxobolus parviformis* (Kallert et al., 2005). However, these three species currently lack genomic resources; only a transcriptome is available only for *M. cerebralis*. Thus, *Myxobolus sp.* represents a promising model to study infection and host-parasite interactions within *Myxobolus* – since both hosts have been molecularly confirmed, a good quality long-read genome has been sequenced, and a transcriptome is available. Further, the maintenance of this life cycle involving fathead minnows in a laboratory setting is much easier and cost-effective than doing the same with salmonid fish.

## 5. Conclusion

We sequenced, assembled, and annotated the genome of *Myxobolus sp.*, an emerging parasite of fathead minnows in Alberta, using the long read Oxford Nanopore sequencing technology. The *Myxobolus sp.* genome is the largest genome among myxozoan species sequenced till date, and one of the most repetitive genomes among animals. We compare the *Myxobolus sp.* genome to genomes from other myxozoan and free-living cnidarian species, and show that, like other myxozoan species, *Myxobolus sp.* has lost several well-conserved eukaryotic genes, and at the same time, has also experienced expansion in several gene families. Our analysis demonstrated the value of long read sequencing platforms for all myxozoan parasites. We also propose that *Myxobolus sp.* presents a promising potential model to study host-parasite interactions in myxozoans.

## Supporting information

Supplemental Results and Methods

## 6. Acknowledgements

We are grateful to Molly Tilley for help collecting *Myxobolus sp.* samples. We acknowledge the high-performance computing resources by the Faculty of Veterinary Medicine and Research Computing at the University of Calgary.

