## Supplemental Results and Methods for "The highly repetitive genome of *Myxobolus* sp., a myxozoan parasite of fathead minnows"

### Supplementary Results

#### 1. Comparing the genome annotations of *M. rasmusseni* using BRAKER with RNAseq and protein data and BRAKER using only protein data

We tried two different BRAKER pipelines to annotate the genome of *M. rasmusseni*.

For BRAKER with RNAseq and protein data, we used the STAR aligner v 2.7.10b_alpha_230301 (Dobin et al., 2013) to map the RNA-seq reads to the *M. rasmusseni* genome and used the resulting mapping file as input for BRAKER. We used the OrthoDB v10 metazoa protein database as protein evidence for BRAKER. The Transcript Selector for BRAKER (TSEBRA) tool was used to combine the BRAKER predictions from both the RNAseq and protein evidence analyses. For BRAKER with protein data alone, the procedure is outlined in Methods and Materials Section 2.4. We used BUSCO on the set of predicted proteins from both BRAKER runs (Table S1). While BRAKER (RNAseq + protein) had more complete BUSCO proteins, the duplication rate was high, compared to BRAKER (protein alone) and to the only other long-read assembly in myxozoans (*M. honghuensis*). We also analyzed the average gene, intron, and exon lengths for the two BRAKER analyses for *M. rasmusseni* and found that when we used only protein evidence alone, the average gene lengths and intron lengths are much lower than when RNAseq and protein evidence both were used and were similar to those reported for *M. honghuensis* (Table S2). Hence, we decided to proceed with genome annotations obtained using BRAKER with protein data alone.

**Table S1**

| | BUSCO on the predicted protein set (Eukaryota database) |
|---|---|
| *M. honghuensis* | C:51.0% [S:49.4%, D:1.6%], F:14.1%, M:34.9% |
| *M. rasmusseni (RNAseq + protein)* | C:47.5% [S:37.3%, D:10.2%], F:13.3%, M:39.2% |
| *M. rasmusseni (only protein)* | C:44.3% [S:41.6%, D:2.7%], F:16.5%, M:39.2% |

**Table S2**

| | Average gene-length | Average intron length | Average exon length |
|---|---|---|---|
| *M. honghuensis* | 3295.028705 | 697.9894182 | 245.9979515 |
| *M. rasmusseni (RNAseq + protein)* | 7717.355008 | 1836.070391 | 250.0092666 |
| *M. rasmusseni (only protein)* | 3357.522759 | 746.6225414 | 267.414841 |

#### 2. Description of the *M. rasmusseni* transcriptome

After filtering the reads generated using Illumina sequencing using Trimmomatic (Bolger et al., 2014), 340,637,040 paired-end reads remained and were assembled into 173,722 transcripts by rnaSPADES (Bushmanova et al., 2019). TransDecoder (Haas, BJ. https://github.com/TransDecoder/TransDecoder) predicted 65,820 coding regions in these transcripts. We used CD-HIT (Fu et al., 2012) to cluster the protein sequences at 100% similarity. The final set of predicted proteins from the *M. rasmusseni* transcriptome was comprised of 54,665 proteins. Next, we removed host contamination from these predicted proteins. We used BLASTP to align these proteins against proteins from the fathead minnow (e-value: 1e-5).

#### 3. Effect of gene-prediction tool on BUSCO analysis in myxozoans

We used BUSCO with both Augustus and MetaEuk options to assess genome completeness using BUSCO (Eukaryota database (255 genes) in genome mode). We used BUSCO with Augustus using the default species (fly) and also the cnidarian *Hydra vulgaris*. We found that for all myxozoan species MetaEuk consistently detected more BUSCO genes compared to Augustus (Figure S1, Table S3).

**Figure S1** The BUSCO analysis of the myxozoan genomes, where (A) refers to Augustus, (A+H) refers to Augustus using *Hydra vulgaris* as the species, and (M) refers to MetaEuk.

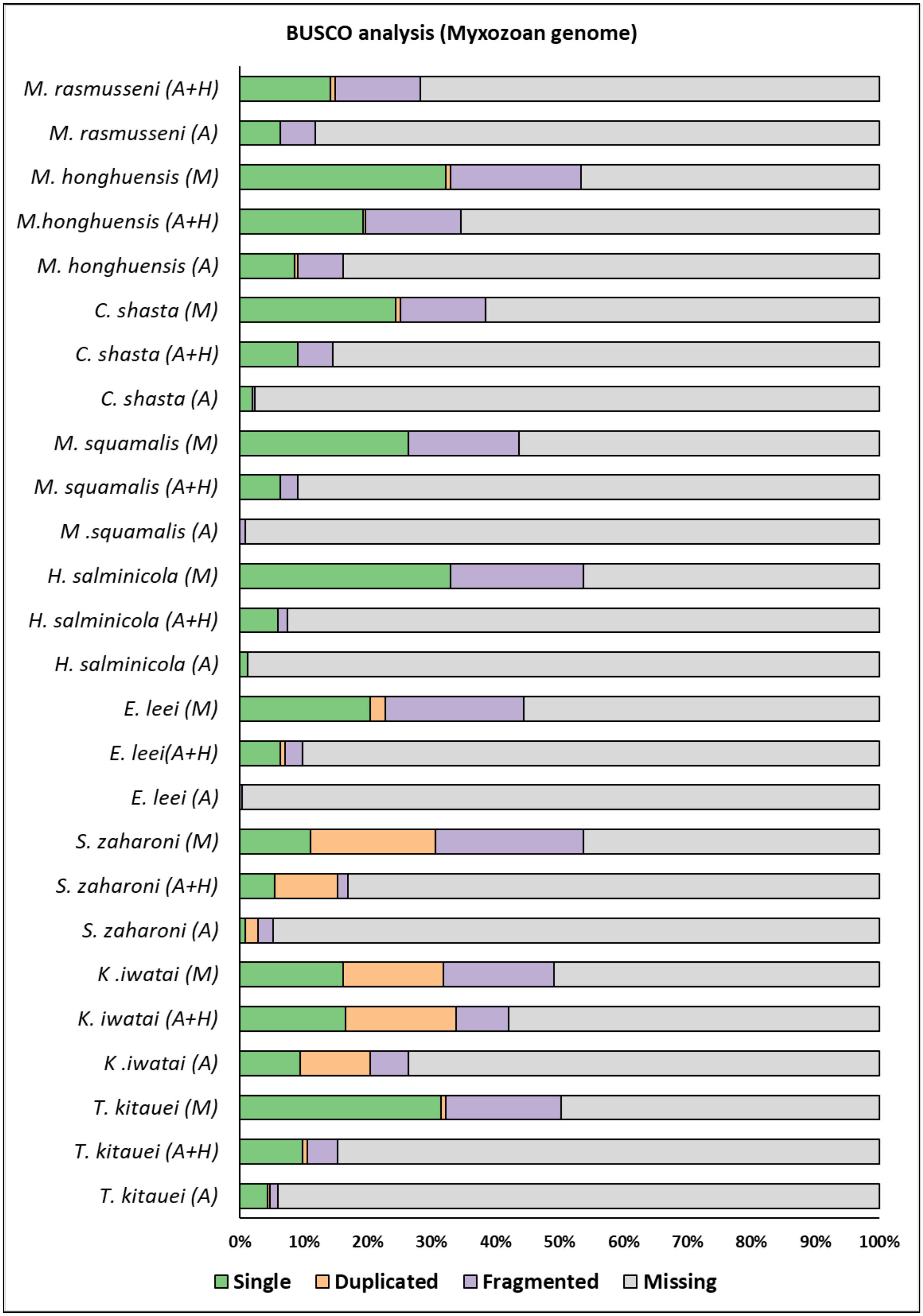
BUSCO analysis (Myxozoan genome)
M. rasmusseni (A+H)
M. rasmusseni (A)
M. honghuensis (M)
M.honghuensis (A+H)
M. honghuensis (A)
C. shasta (M)
C. shasta (A+H)
C. shasta (A)
M. squamalis (M)
M. squamalis (A+H)
M .squamalis (A)
H. salminicola (M)
H. salminicola (A+H)
H. salminicola (A)
E. leei (M)
E. leei(A+H)
E. leei (A)
S. zaharoni (M)
S. zaharoni (A+H)
S. zaharoni (A)
K .iwatai (M)
K. iwatai (A+H)
K .iwatai (A)
T. kitauei (M)
T. kitauei (A+H)
T. kitauei (A)
0% 10% 20% 30% 40% 50% 60% 70% 80% 90% 100%
Single Duplicated Fragmented Missing

**Table S3** The BUSCO analysis of the myxozoan genomes, where (A) refers to Augustus, (A+H) refers to Augustus using *Hydra vulgaris* as the species, and (M) refers to MetaEuk.

| Species | Single | Duplicate d | Fragmented | Missing |
|---|---|---|---|---|
| *T. kitauei (A)* | 4.3% | 0.4% | 1.2% | 94.1% |
| *T. kitauei (A+H)* | 9.8% | 0.8% | 4.7% | 84.7% |
| *T. kitauei (M)* | 31.4% | 0.8% | 18.0% | 49.8% |
| *K .iwatai (A)* | 9.4% | 11.0% | 5.9% | 73.7% |
| *K. iwatai (A+H)* | 16.5% | 17.3% | 8.2% | 58.0% |
| *K .iwatai (M)* | 16.1% | 15.7% | 17.3% | 50.9% |
| *S. zaharoni (A)* | 0.8% | 2.0% | 2.4% | 94.8% |
| *S. zaharoni (A+H)* | 5.5% | 9.8% | 1.6% | 83.1% |
| *S. zaharoni (M)* | 11.0% | 19.6% | 23.1% | 46.3% |
| *E. leei (A)* | 0.0% | 0.0% | 0.4% | 99.6% |
| *E. leei(A+H)* | 6.3% | 0.8% | 2.7% | 90.2% |
| *E. leei (M)* | 20.4% | 2.4% | 21.6% | 55.6% |
| *H. salminicola (A)* | 1.2% | 0.0% | 0.0% | 98.8% |
| *H. salminicola (A+H)* | 5.9% | 0.0% | 1.6% | 92.5% |
| *H. salminicola (M)* | 32.9% | 0.0% | 20.8% | 46.3% |
| *M .squamalis (A)* | 0.0% | 0.0% | 0.8% | 99.2% |
| *M. squamalis (A+H)* | 6.3% | 0.0% | 2.7% | 91.0% |
| *M. squamalis (M)* | 26.3% | 0.0% | 17.3% | 56.4% |
| *C. shasta (A)* | 2.0% | 0.0% | 0.4% | 97.6% |
| *C. shasta (A+H)* | 9.0% | 0.0% | 5.5% | 85.5% |
| *C. shasta (M)* | 24.3% | 0.8% | 13.3% | 61.6% |

| | | | | |
|---|---|---|---|---|
| *M. honghuensis (A)* | 8.6% | 0.4% | 7.1% | 83.9% |
| *M.honghuensis (A+H)* | 19.2% | 0.4% | 14.9% | 65.5% |
| *M. honghuensis (M)* | 32.2% | 0.8% | 20.4% | 46.6% |
| *M. rasmusseni (A)* | 6.3% | 0.0% | 5.5% | 88.2% |
| *M. rasmusseni (A+H)* | 14.1% | 0.8% | 13.3% | 71.8% |
| *M. rasmusseni (M)* | 31.0% | 0.4% | 19.2% | 49.4% |

#### Supplementary File S1

Here, we list the steps used to extract DNA from myxospores.

### HMW DNA for PacBio

1. **EXTRACTION** MagAttract® HMW DNA Qiagen kit, Ref.67563

You will need:

- A magnetic rack
- A rotator

It requires 4 days:

- Day 1: clean up (if ethanol preserved) and overnight digestion
- Day 2: buffers for extraction
- Day 3: first DNA collection
- Day 4: second DNA collection (and clean-up)

If samples are on EtOH, they should be cleaned as follow:

- On a platform rocker or tube rotator, incubate your sample in an ample volume of Buffer STE (400 mM NaCl, 20 mM Tris pH 7.5, 30 mM EDTA) for 10-15 minutes.
- Replace the solution with the same volume of fresh STE and incubate on a platform rocker or tube rotator another 10-15 minutes.
- Transfer the tissue to a chilled surface (such as a clean aluminum block in an ice bucket) and wick the excess liquid with a Kim Wipe.
- Proceed with the DNA extraction.

If samples were snap frozen right after collection, and kept at -80C, you can start directly with extraction

1. Add 220 µl Buffer ATL to the sample.
2. Add 20 µl Proteinase K and mix by vortexing.
3. Incubate overnight (12–16 h) at 56°C, shaking at 900 rpm until the tissue is completely lysed.

Spin down the tube at maximum speed to remove droplets from the lid and to ensure that the incompletely lysed tissue particles are settled at the bottom of the tube.

4. Transfer 200 µl of the lysate to a new 2 ml sample tube.
5. Add 4 µl RNase A to the sample. Mix by pulse-vortexing and incubate for 2 min at room temperature
6. Add 150 µl Buffer AL to the sample. Mix by pipetting up and down.
7. Add 280 µl Buffer MB to the sample.
8. Add 40 µl of MagAttract Suspension G to the sample.
9. Place the tube in the rotator and allow for binding at room temperature for 15 min.
10. Place the tube onto the magnetic base, wait until bead separation has been completed (~1 min), and remove all the supernatant without disturbing the magnetic bead pellet.
11. Remove tube from magnetic base and add 700 µl Buffer MW1 directly onto the magnetic bead pellet and place in rotator for 10 minutes ($2^{nd}$ wash for 5 min)
    a. Place the tube onto the magnetic rack, wait until bead separation has been completed (~1 min), and remove the supernatant.
12. Repeat 11 and 11.a
13. Remove tube from magnetic base and add 700 µl Buffer PE directly onto the magnetic bead pellet and place in rotator for 10 minutes ($2^{nd}$ wash for 5 min)
    a. Place the tube onto the magnetic rack, wait until bead separation has been completed (~1 min), and remove the supernatant.
14. Repeat steps 13 and 13.a

**Note:** Remove all the supernatant. Use a small pipette tip to remove any traces of Buffer PE.

15. Rinse the particles with 700 µl distilled water while the tube is on the magnetic base and the beads are fixed to the wall of the sample tube. Incubate for 1 min at room temperature and remove the supernatant.

**Important:** Do not pipet water directly onto the bead pellet – pipet it into the sample tube against the side facing away from the bead pellet.

16. Repeat step 15.
17. Remove the tube from magnetic base and add 20 µl of Buffer AE (preheated at 55-60°C). Place the tube in the rotator and allow for binding at room temperature for 15 min.
18. Quick spin to get everything to the bottom of the tube and let it sit overnight.
19. Place the tube on the magnetic base, wait until bead separation has been completed (~1 min), and transfer the supernatant with the high-molecular-weight DNA to a new sample tube.
20. Repeat step 17 (second time with 15 uL of buffer AE), 18 and 19.

2. **CLEAN-UP** This step is performed using a combination of 2 kits, we use the beads from Select-a-size Magbead kit, Cat# D4084-3-10 and the buffers from Genomic DNA Clean&Concentrator -25, Cat# D4064, both from Zymo Research.

You will need:

- A magnetic rack
- A rotator

1. Add 4 volumes of DNA Binding Buffer to each sample and mix well.
2. Add 16.5 µl MagBinding Beads and mix well for 15-20 minutes (pippette constantly for suspension)

Important: MagBinding Beads settle quickly, ensure that beads are kept in suspension while dispensing.

3. Transfer the tube to the magnetic stand until beads have pelleted, then aspirate and discard the cleared supernatant.
4. Add 500 ul of DNA Wash Buffer and mix well. Pellet the beads and discard the supernatant.
5. Repeat Step 4.
6. Dry the beads at room temperature for 10 minutes or until dry (beads should go from glossy to matte).
7. To elute DNA from the beads, add 25 ul DNA Elution Buffer (heated up to 60C) and mix well for 5 minutes.
8. Transfer the tube to the magnetic stand until beads have pelleted, then aspirate and dispense the eluted RNA to a new tube.

The eluted DNA can be used immediately or stored frozen.
